# Drug-tolerant persister B-cell precursor acute lymphoblastic leukemia cells

**DOI:** 10.1101/2023.02.28.530540

**Authors:** Mingfeng Zhang, Lu Yang, David Chen, Nora Heisterkamp

## Abstract

Reduced responsiveness of precursor B-acute lymphoblastic leukemia (BCP-ALL) to chemotherapy can be inferred when leukemia cells persist after 28 days of initial treatment. Survival of these long-term persister (LTP) / minimal residual disease (MRD) cells is partly due to bone marrow stromal cells that protect them under conditions of chemotherapy stress. We used RNA-seq to analyse BCP-ALL cells that survived a long-term, 30-day vincristine chemotherapy treatment while in co-culture with bone marrow stromal cells. RNAs of as many as 10% of the protein-encoding genes were differentially expressed. There was substantial overlap with genes associated with MRD cell persistence reported in other studies. The top pathway regulated in the LTP cells was that involving p53, a master regulator of a spectrum of responses relevant to drug resistance and cytotoxic drug exposure including control of autophagy. We tested a select number of genes for contribution to BCP-ALL cell survival using Cas9/CRISPR in a 2-step selection, initially for overall effect on cell fitness, followed by 21 days of exposure to vincristine. Many genes involved in autophagy and lysosomal function were found to contribute to survival both at steady-state and during drug treatment. We also identified MYH9, NCSTN and KIAA2013 as specific genes contributing to fitness of BCP-ALL cells. CD44 was not essential for growth under steady state conditions but was needed for survival of vincristine treatment. Finally, although the drug transporter ABCC1/MRP1 is not overexpressed in BCP-ALL, a functional gene was needed for DTP cells to survive treatment with vincristine. This suggests that addition of possible ABCC1 inhibitors during induction therapy could provide benefit in eradication of minimal residual disease in patients treated with a chemotherapy regimen that includes vincristine.

## Introduction

Precursor B-cell acute lymphoblastic leukemia (BCP-ALL) develops in the bone marrow after malignant transformation of normal early B-lineage hematopoietic cell precursors. BCP-ALL is one of the most common types of childhood cancers worldwide but presenting overall long-term survival rates of close to 90% in this age-group. In adult patients however, treatment success rates are drastically lower, with only 30-40% achieving long-term remission. Standard initial therapy of BCP-ALL typically consists of a combination of a chemotherapeutic agent (*e.g*. vincristine), a glucocorticoid (*e.g*. prednisone) and an asparaginase (1, 2). Vincristine is a vinca alkaloid chemotherapeutic agent used at several stages of BCP-ALL treatment. This compound interacts with the β-tubulin subunit of α/β-tubulin to disrupt the polymerization of microtubules, which arrests the cell cycle at the M phase leading to apoptosis (3).

The first-line treatment typically encompasses a period of so-called induction therapy in which the patient is treated for 28 days with chemotherapy to induce a remission. Induction therapy usually is able to drastically reduce the number of leukemic cells. A bone marrow sample is subsequently taken to determine the number of remaining leukemia cells, which are called minimal residual disease (MDR). MRD is a key prognostic factor, with higher MRD levels predicting worse outcomes for patients (4). Chemoresistance, as detected by higher MRD levels, thus remains a major clinical problem (5). MRD persists not only by leukemia cell-intrinsic mechanisms such as clonal selection and genetic changes, but also through protective interactions of the leukemia cells with the surrounding supporting stroma. This mechanism of drug resistance has been named environment-mediated drug resistance (EMDR) and is a general route through which MRD cells, regardless of underlying genetics of the leukemia cells, can persist after different types of chemotherapy (6, 7).

EMDR can be studied in tissue culture models by treating leukemia cells with a non-lethal dose of a drug in the presence of stromal cell protection. In the current study, we co-cultured human BCP-ALL cells with mitotically inactivated supporting stromal cells using a long-term drug treatment protocol to investigate factors contributing to the survival of such drug-tolerant persister (DTP) cells. We previously identified specific carbohydrate-binding proteins Galectin-1 and Galectin-3 as contributing to the emergence of DTP leukemia cells (8−10). We here used a general approach including RNAseq and a Cas9/CRISPR dropout screen to investigate other genes that could promote survival of human BCP-ALL cells growing in the long-term presence of vincristine while receiving stromal support.

## EXPERIMENTAL PROCEDURES

### Cell culture and drugs

ICN13 PDX/patient-derived BCP-ALL cells were previously described. They were originally derived from a 15 year-old female leukemia patient with an *MLL-AF4* t(4;11)(q21;q23) translocation (11, 12). ICN13 cells are stromal-dependent and are grown in co-culture with mitotically inactivated OP9 bone marrow stromal cells (ATCC CRL-2749). They were STR genotyped to confirm their identity. To prepare OP9 cells for co-culture, they were allowed to adhere overnight, then treated with 10 μg/ml mitomycin C (Sigma Cat#M4287) for 3 hrs in complete medium, washed, and used. Co-culture was in α-MEM media supplemented with 20% FBS, 1% L-glutamine and 1% penicillin/streptomycin (Life Technologies, Grand Island, NY). A vincristine sulfate solution was obtained from Hospira Worldwide Inc. (Lake Forest, IL, USA). Vincristine sulphate diluted in PBS at different concentrations was added freshly every 3 days. Cells cultured with 2 nM or 4 nM vincristine were collected from triplicate wells at days 18 and 30, respectively, along with controls cultured for the same period of time but without drug (PBS controls). Each biological replicate was divided into two fractions of 2×10^6^ cells, one for proteomic, glycoproteomic and glycomic analyses, and the second for RNA sequencing. Samples were stored at −80°C until analysis. Results of the proteomic, glycomic and glycoproteomic analysis will be the focus of a separate report.

### RNA-seq and expression analysis

RNA-seq analysis was performed as previously described (13). Significantly regulated protein-encoding genes [10,342 total] were defined as fold change ≥2; p<0.05 and low expression filter set at rpkm <1.0. Using these criteria, 276 genes had differential expression in the d18 2 nM vincristine treated group [182 increased and 94 decreased compared to PBS controls]. In the d30, 4 nM treatment group 946 genes were differentially expressed [with 575 increased and 371 decreased]. When both treatment results were combined for statistical analysis, 445 genes met the stated criteria [with 331 showing increased expression and 114 decreased expression compared to PBS control]. Graphs showing normalized RNA counts were generated using GraphPad Prism (v8.4.3). QIAGEN Ingenuity Pathway Analysis (IPA) version 62089861 was used to analyse results of RNA-seq for the d30/4 nM vincristine data set for pathways with differential regulation using p-values and FDR at <0.05 and logFc at - 1.0 – 1.0, resulting in 881 genes to be included in the analysis. RNA-seq data were deposited in GEO under accession number GSE176366.

MRD199 from GSE83142 [Table S7 in Ebinger *et al* (14)] was a constitutional trisomy 21, acquired homozygous 9p deleted BCP-ALL sample. For the comparison, only the 10,167 protein-encoding genes were included, of which 115 were lower and 557 higher expressed in the MRD199 cells compared to BCP-ALL cells from non-drug treated mice. For the patient MRD samples, of the 10,448 protein-encoding genes 334 genes were reduced and 533 increased in expression compared to the diagnosis samples. Raw RNAseq data of 10 matched diagnosis/relapse BCP-ALL samples were downloaded from the NCBI sequence read archive SRA048657. Reads were quality-checked and processed as previously described in detail (13).

### Cas9-CRISPR screen

We used KOPN1 MLL-r B-ALL cells stably transduced with a Cas9-encoding lentivirus as described (13). KOPN1 cells do not need stromal support for steady-state growth in tissue culture but were plated on OP9 cells after selection with blasticidin. Controls included neutral selection genes [neg, LacZ, Luc and Ren] covered by ten sgRNAs each. Essential gene controls were represented by 2 sgRNAs each, including BRD4, CDK9, MYC, PCNA, POLR2A, POLR2D, PRL9, RPA3, RPL2 and RPS20. The 97 target genes were each covered by 10 sgRNAs. In total approximately 1000 sgRNAs were included in the library. Data analysis using model-based analysis of genome-wide CRISPR-Cas9 knockout (MAGeCK, (15, 16)) was as previously described (13) using input n=2; overall survival (n=4 samples) and 1 nM vincristine treatment (n=4) samples. Neg | rank is the ranking of the particular gene in negative selection; neg | score is the robust rank aggregation (RRA) low value of the particular gene in negative selection.

## RESULTS

### Transcriptome

An *ex vivo* co-culture with OP9 bone marrow stromal cells is widely used to support survival and growth of human BCP-ALL cells (8–10, 17–21) and can be used to monitor drug treatment outcomes. In this system, leukemia cells are motile, migrating back and forth underneath the stromal cells (**Fig. 1A**). We treated co-cultures with different concentrations of vincristine over an extended period of time (18 or 30 days, d18 and d30 respectively) to generate DTP leukemia cells that can grow in the continued presence of chemotherapy (**Fig. 1B, C**) as a model for MRD in patients after 30 days of induction chemotherapy.

**Figure 1.**
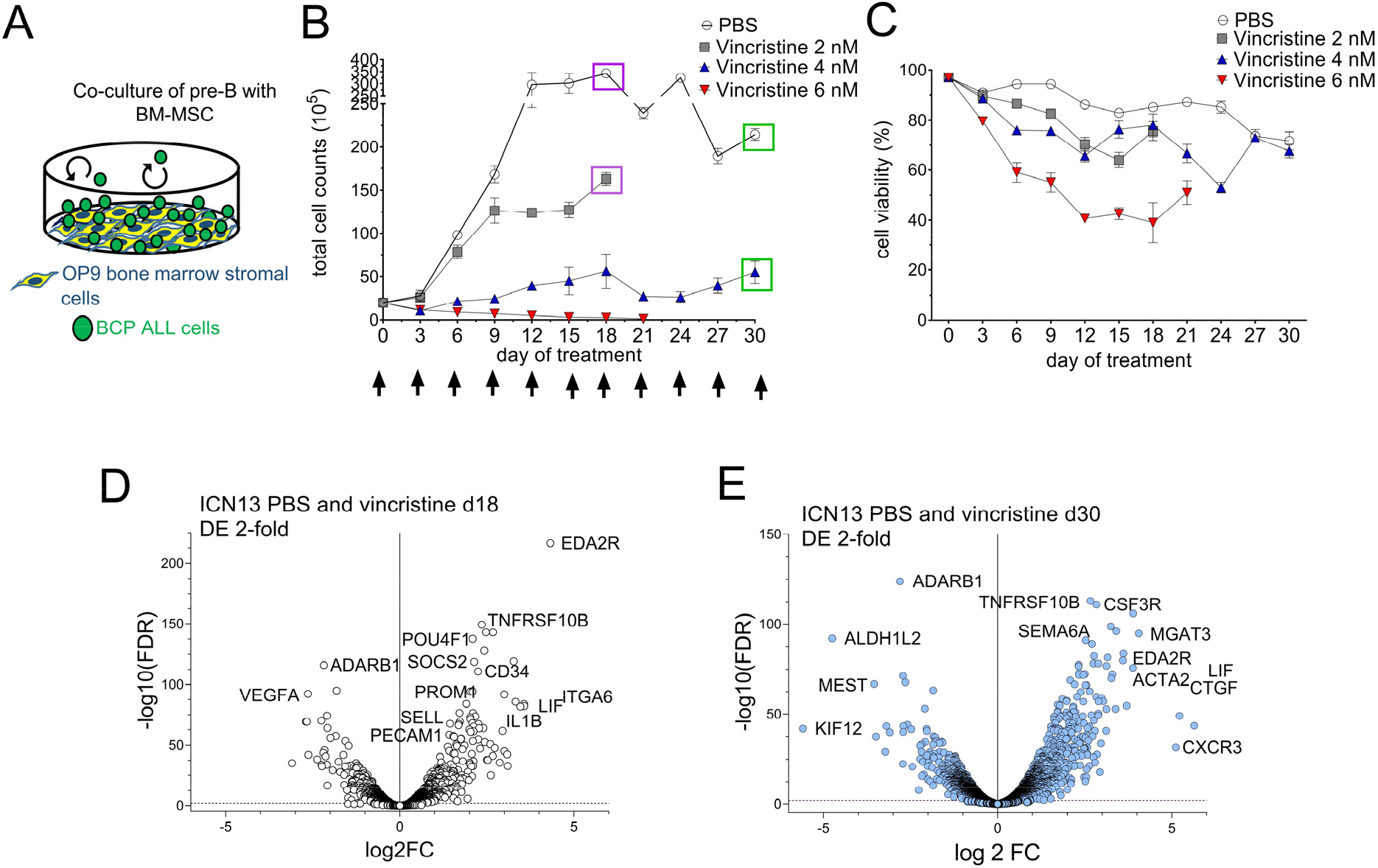
Analysis of drug insensitivity development of human BCP-ALL cells treated with the chemotherapy drug vincristine. (**A**) Schematic illustration of co-culture of leukemia cells with stromal support. Leukemia cells traffic dynamically between the stromal support layer and the medium; cells were harvested from the supernatant for omics analysis on the days indicated by purple and green boxes in (**B**). (**B, C**) Total cell numbers and viability (defined as number of viable cells/total cell number x 100) of the BCP-ALL cells. Arrows in panel B indicate that fresh vincristine was added every third day. Viable cell counts were performed using Trypan blue exclusion. (**D**) Volcano plot representing 182 genes with increased and 94 genes with decreased expression on d18 of 2 nM vincristine treatment compared to PBS controls grown in parallel cultures (**Table S1 T2**). (**E**) Volcano plot depicting 575 genes with increased transcript levels and 372 down-regulated transcripts on d30 of 4 nM vincristine treatment. Each point represents the mean of biological triplicate samples. Also see **Table S1 T2**).

The effect of this cytotoxic drug on the leukemia cells is dose-dependent: after 6 days of 2 nM chemotherapy, cells were able to resume proliferation, and their viability was relatively good compared to the PBS controls. Cells treated with 4 nM grew very slowly whereas 6 nM vincristine completely inhibited proliferation (**Fig. 1B**). Therefore, we harvested cells for RNA-seq (**Fig. 1B**) from the 2 and 4 nM treated cultures when sufficient cells had accumulated, on d18 and d30 of culture, respectively. Analysis of the results showed that protein-encoding transcripts of just 276 genes showed significant differential expression (DE) after 18 days of treatment with 2 nM vincristine (**Fig. 1D, TableS1 Tab 1).** Exposing cells to the higher drug dose (4 nM) over a longer period of 30 days resulted in a differential expression of around 946 genes (**Fig. 1E, Table S1, Tab 2**).

### DTP-MRD-associated genes

Two studies have compared gene expression of actively growing BCP-ALL cells to that of leukemia cells treated with chemotherapy within a time frame used clinically for induction chemotherapy. Ebinger *et al* (14) compared MRD cells from patients treated with induction chemotherapy to diagnosis samples before the start of treatment, and also from mice transplanted with BCP-ALL cells and treated for 14 days with vincristine and cyclophosphamide. We next determined if there was overlap between the genes with DE in our study and the MRD genes reported by Ebinger et al. **Figure S1A** (**Table S1 Tab 4**) shows that 146 genes with DE in ICN13 cells growing at d30 in 2 nM vincristine also had DE in MRD cells recovered from mice transplanted with BCP-ALL-199 cells after 14 days of chemotherapy, of which 113 (77%) were regulated in the same direction. There was also a substantial overlap of 110 DE genes between patient MRD and ICN13 d30 drug treated samples (**Figure S1B** and **Table S1 tab T5**) of which 68 (62%) were concordant in direction. Finally, there were 21 genes with common regulation in all three MRD data sets (**Table S1 tab T6**). 13 of these genes also were represented in and had differential expression in GSE67684 [Yeoh *et al* (22)] containing microarray gene profiling data of 220 pediatric BCP-ALL patient samples on d0, d8, d15 and d33 of chemotherapy (**Figure S1C** and **Table S1 tab T7**). This analysis indicates that the transcriptome of DTP cells generated in a tissue coculture model has common features with MRD leukemia persisting after standard chemotherapy treatment.

We also compared differentially expressed genes in ICN13 at d30 of vincristine treatment to differentially expressed genes in BCP-ALL relapsed samples compared to diagnosis samples (23), finding 38 genes in common, of which 28 (74%) were regulated in the same direction (**Figure S1D** and **Table S1 Tab 8**). However, this set of genes did not overlap with the 21 common DTP/MRD genes suggesting a fundamental difference between cells that survive induction chemotherapy and cells from relapsed patients..

### GSEA

We used the ICN13 d30 4 nM vincristine treatment results to further investigate if changes in gene expression in the *in vitro* generated MRD cells point to significantly modified pathways using gene set enrichment analysis (GSEA (24)). Not surprisingly, the top down regulated gene sets belonged to the G2M checkpoint, E2F pathway (25) and mitotic spindle pathways, consistent with the effects of attenuation of cell cycle via microtubule polymerization through vincristine. Interestingly, based on top enrichment scores, among others, coagulation, IL6/JAK/STAT3 and complement pathways (**Figure S2A**, **Table S1 tab T9**) were activated.

We previously analysed changes in gene expression of murine BCP-ALL cells developing drug insensitivity when co-cultured with mouse embryonic fibroblasts as feeder layer and treated over a period of up to 30 days (20). Murine leukemia cells were Bcr/Abl-positive and drugs included the farnesyltransferase inhibitor lonafarnib and the tyrosine kinase inhibitor nilotinib (20). We performed GSEA on these data for comparison with the results of the human ICN13 BCP-ALL cells treated with vincristine. As illustrated in **Figure S2B (Table S1 tab T10)**, even though very different drugs as well as subtypes of BCP-ALL were used, there was a surprising concordance between pathways altered in these two data sets. This included decreased activation of E2F targets and G2M checkpoints and upregulation of inflammatory responses, IL6/JAK/STAT3 signalling pathways, complement and coagulation pathways. p53 involvement in the mouse LTP cells was not found, perhaps because of the differences between murine and human p53 function (26).

We also used QIAGEN Ingenuity pathway analysis (IPA) to further examine differentially expressed genes in the ICN13 d30 vincristine data set. As shown in **Figure 2** (**Table S1 tab T11**), numerous canonical pathways were significantly regulated when comparing the d30 vincristine-treated BCP-ALL cells with their non-drug treated controls. Consistent with GSEA, kinetochore metaphase signalling was decreased as expected (**Table S1, Tab T16**). Pathways with differential gene expression consistent with activation included integrins, thrombin (**Figure S3**) and HMBG1 (**Figure S4**) (**Table S1, Tab T13-15**). Other interesting patterns detected in this analysis were in G2M checkpoint and inflammatory responses (**Table S1, Tab T17, 18 and 19** respectively).

**Figure 2.**
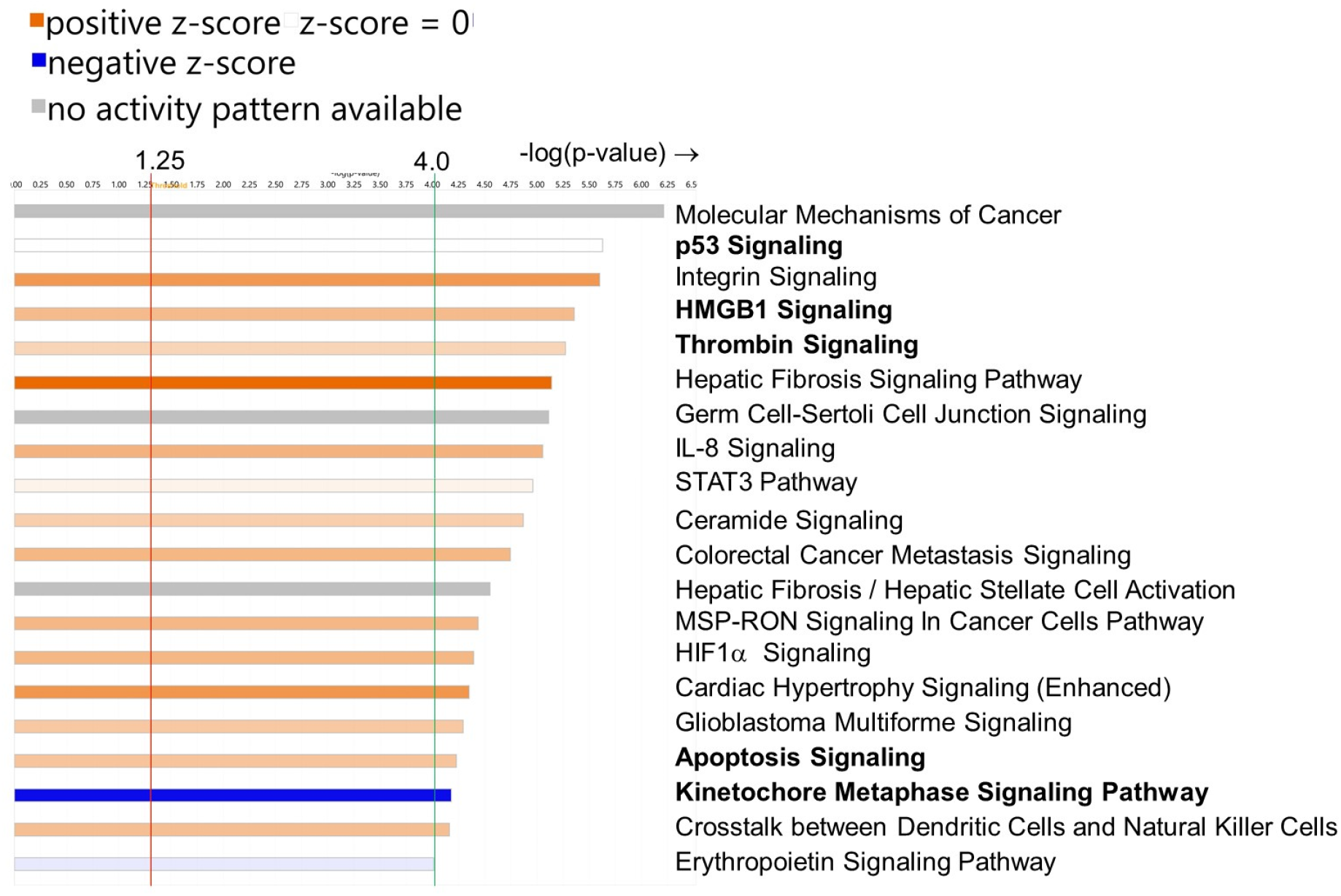
Analysis of canonical pathways significantly altered in ICN13 BCP-ALL cells treated for 30 days with 4 nM vincristine compared to control cells cultured in parallel without vincristine treatment. IPA^®^ analysis included 881 genes with differential expression (FDR and p-value <0.05) and logFc between −1.0 and +1.0. Orange line: threshold at −log(p-value) of 1.25. Green line - pathways with −log(p-value) >4.0 as shown here. Also see **Table S1, Tab 11**.

The most significant and specific canonical pathway emerging from this analysis was that of p53 **(Table S1 tab T12)**. This was also one of the top pathways indicated in the GSEA of long-term drug surviving human ICN13 BCP-ALL treated with vincristine (**Figure S2A**) but not mouse cells. Genes with differential expression in ICN13 cells controlled by this master regulator are involved in a variety of functions (**Figure 3A**) of which cell cycle progression/cell cycle arrest (**Figure 3B**), apoptosis, cell survival, DNA repair, senescence (**Figure 3C, 3D**), glycolysis and autophagy (**Figure 3D**) were significantly altered in the ICN13 DTP cells.

**Figure 3.**
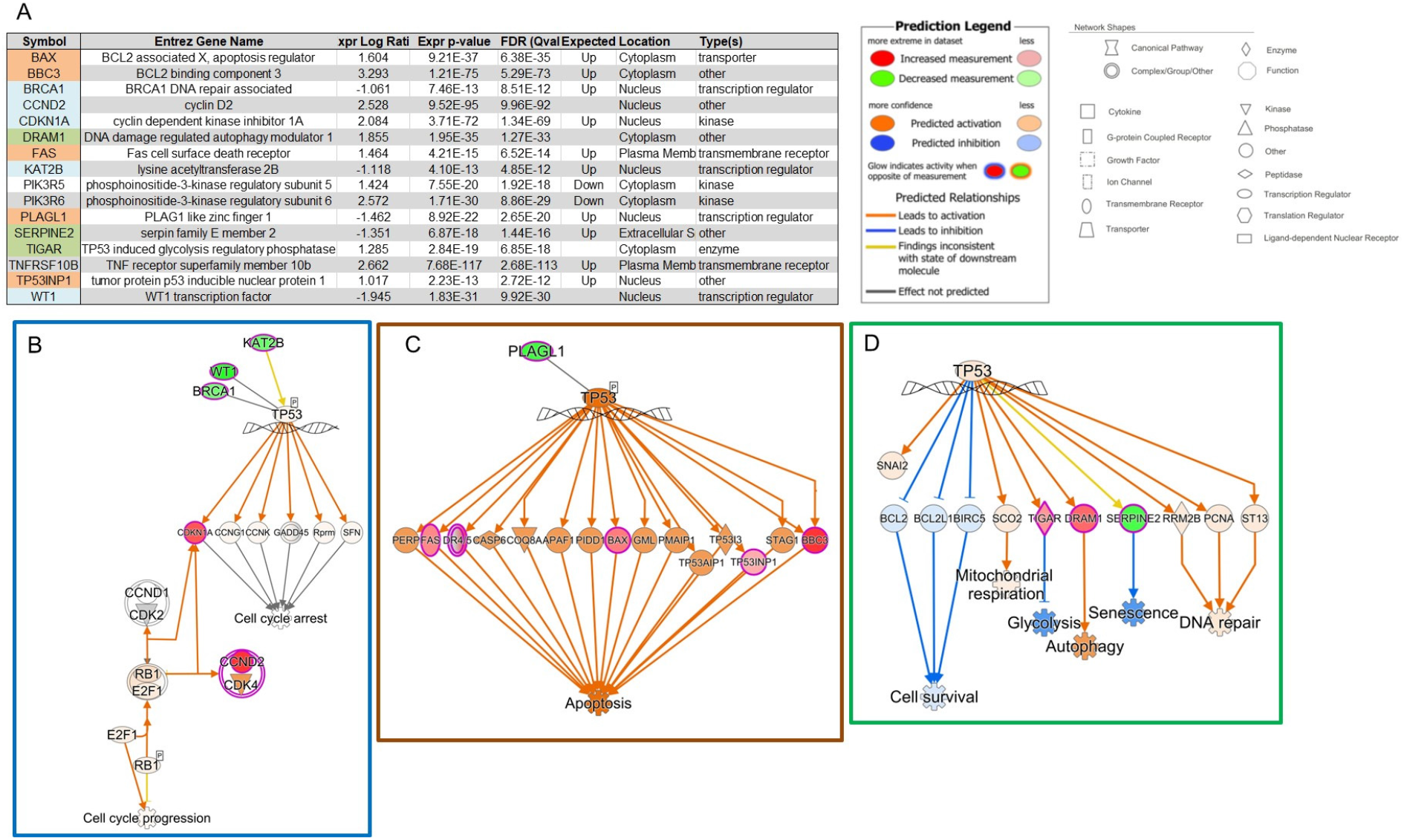
Engagement of different p53-related pathways based on differentially expressed genes in ICN13 BCP-ALL cells treated for 30 days with 4 nM vincristine. IPA^®^ analysis (**A**) Left, details of genes included in this analysis. Right, key to schematic diagrams in B-D. (**B**) Regulation of cell cycle by p53 (**C**) Regulation of apoptosis by p53 (**D**) p53 control of functions associated with other key survival pathways in cancer cells including autophagy through DRAM1.

### Cas9-CRISPR dropout screen

We selected a small number of the DTP/MRD signature genes for Cas9/CRISPR knockout to determine if their overexpression could contribute to the phenotype of these cells (**Table S2 tabs T1-3**). Similarly, differential expression in BCP-ALL patient samples comparing expression on d0, d8, d15 and d33 of induction chemotherapy could indicate a cellular response to mitigate the effects of chemotherapy (**Table S2 tab T4**). Genes with increased expression in BCP-ALL cells compared to normal control CD19+CD10+ cells could also provide a persistence advantage when challenged by chemotherapy (**Figure S5** and **Table S2 tab T5**).

One of the most well-studied mechanisms of general drug resistance development involves the amplification and/or overexpression of drug efflux transporters. Of the 42 ABC family of genes present in humans, only few were reported to function as an exporter for xenobiotic drugs. Based on expression levels in BCP-ALL cells (**Table S2 tab T6**) we selected 4 of the known MDR genes (27) for knockout using Cas9/CRISPR. Of note, none of these showed consistently increased expression in DTP/MRD BCP-ALL cells.

A more recently identified mechanism of general drug resistance is lysosomal sequestration of drugs including among others tyrosine kinase inhibitors and vincristine (28, 29). Based the importance of autophagy in survival of stressed cells [reviewed in (30, 31)] and on the activation of the p53 pathway noted in the ICN13 cells surviving long term vincristine treatment, we included a selection of genes involved in autophagy and lysosomal homeostasis for the Cas9/CRISPR knockout.

In total we selected a limited number of around 100 genes to increase the sensitivity of detection (**Table S2 tab T7**). About 20% of these genes are designated ‘strongly selective’ or ‘common essential’ genes in other cancer cell lines (DEPMAP; CRISPR and RNAi **Table S2 tab T8**). The experimental set-up is schematically shown in **Figure 4A**. We used KOPN1, an MLL-r BCP-ALL cell line constitutively expressing Cas9, for this screen. Cells were harvested on d27 and d48 to first detect loss of genes that contribute to general survival of these cells without the stress of chemotherapy. These samples were compared to input counts. Subsequently cells were co-cultured with OP9 stomal cells and then treated in biological duplicates (**Figure 6 B1** and **B2**) with 1 nM vincristine for 15-21 days. We compared DNA counts of cells before treatment to those at the end of treatment to detect genes specifically needed to promote persistence under vincristine chemotherapy. **Table S3** summarizes the data and analysis results by MAGeCK (16).

**Figure 4.**
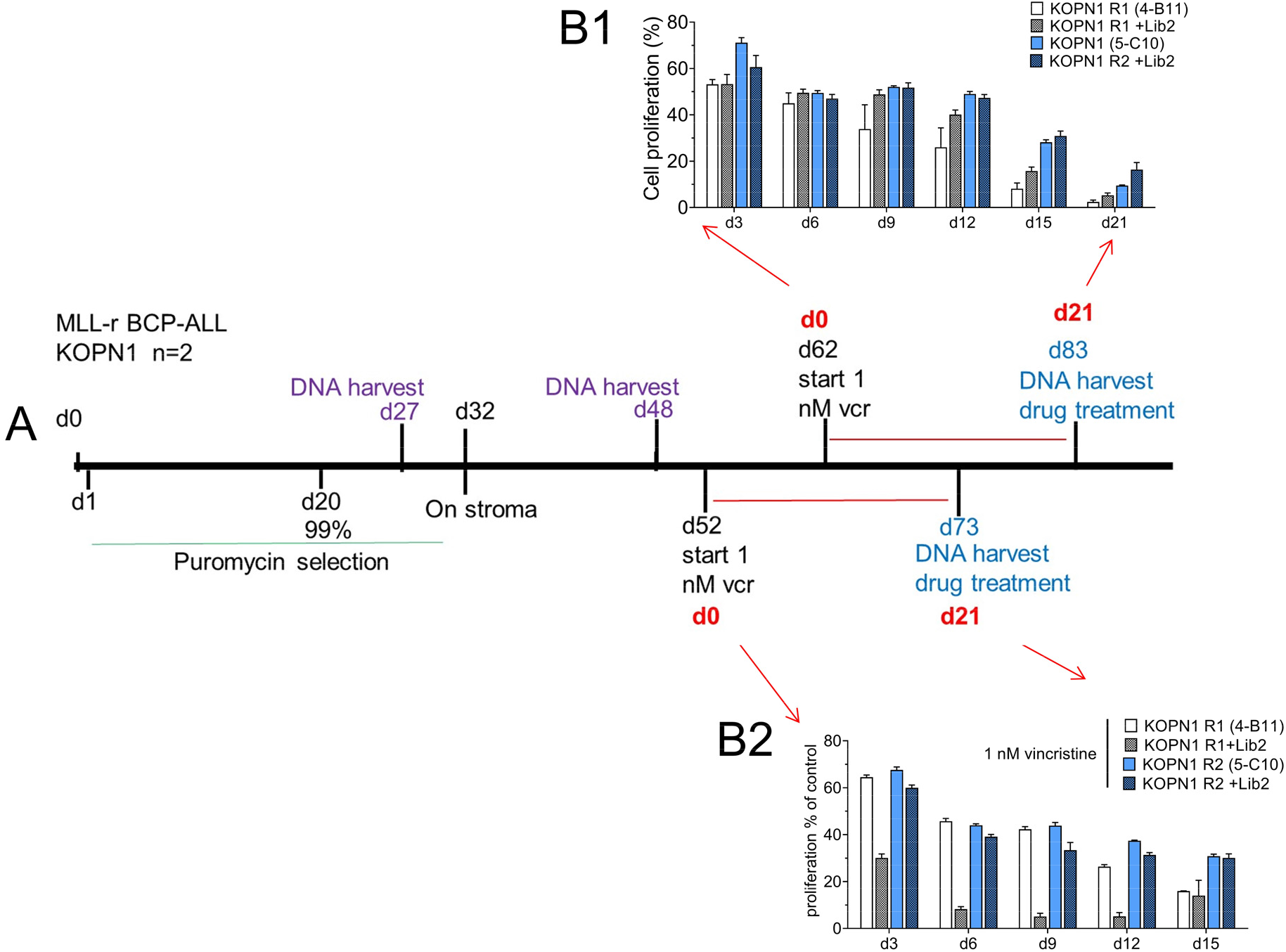
Schematic of Cas9/CRISPR screen on KOPN1 MLL-r BCP-ALL cells. **(A)** Schematic timeline of CRISPR/Cas9 screen targeting 97 selected genes. In brief, transduction of cells with the lentiviral sgRNA library at a low MOI on d0 was followed by puromycin selection to obtain a population in which 99% of cells contained a lentiviral construct. Two different aliquots of KOPN1 (R1, 4-B11 and R2, 5-C10) were independently transduced. For detection of genes needed for cell survival without drug treatment, DNA of transduced cells was harvested at d27 and 48 of selection as indicated for an n=4 samples. Cells were plated on mitotically inactivated OP9 stromal cells at d32 to provide support during vincristine drug treatment. Drug treatment with 1 nM vincristine was applied for 21 days, commencing on d52 or on d62 as indicated. DNA (n=4 total drug-treated samples) was isolated on the indicated days. (**B1, B2**) Growth of the KOPN1 cells transduced with the library (+Lib2) over the period of vincristine treatment compared to parental nontransduced KOPN1 cells (R1, 4-B11 and R2, 5-C10) treated in parallel with 1 nM vincristine. Proliferation was monitored by CellTiterGlo assay for ATP and expressed as a percentage compared to non-drug treated cells grown in parallel.

**Figure 5.**
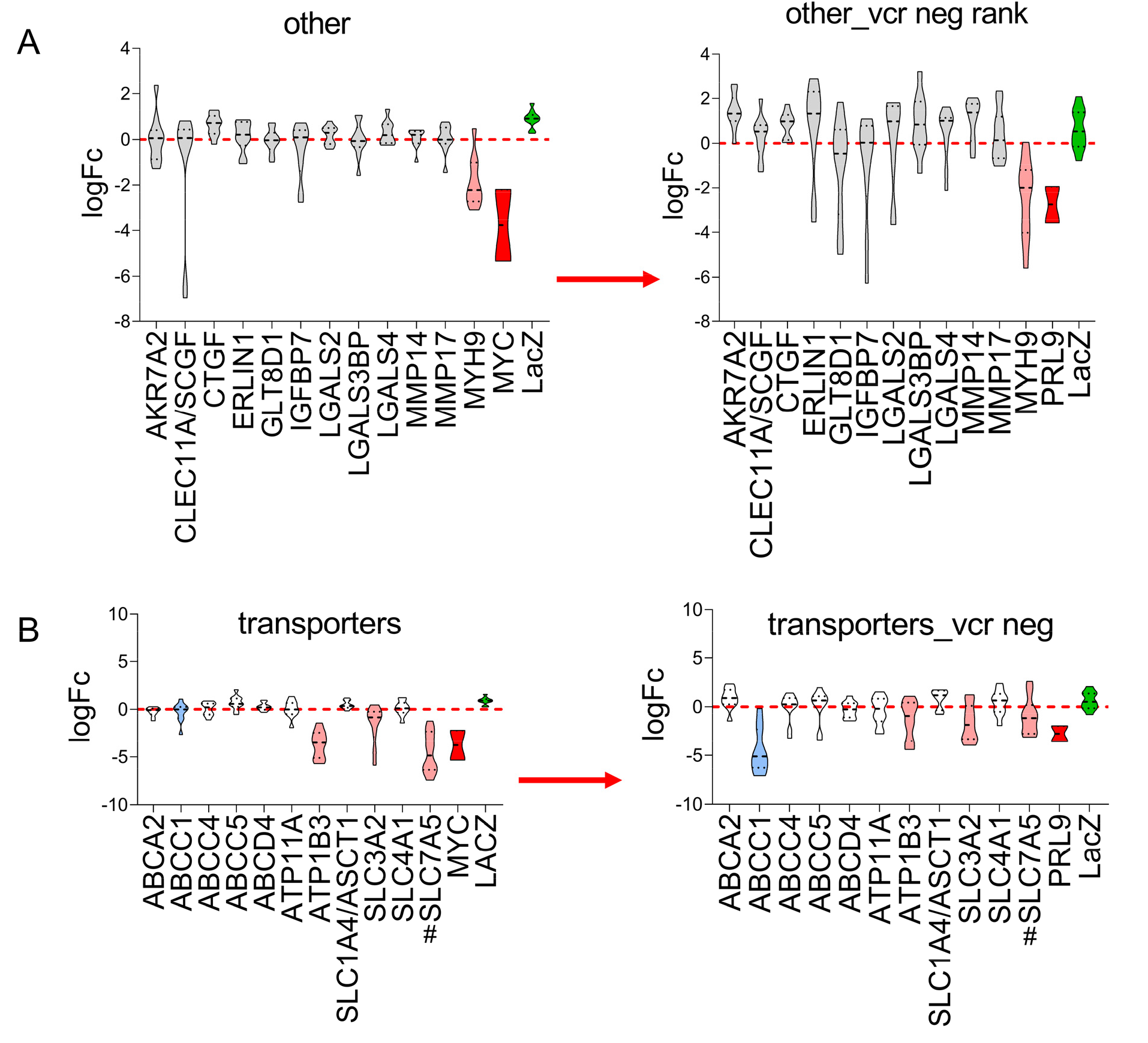
Contribution of selected genes to overall survival and drug-tolerant persister state of KOPN1 BCP-ALL cells. Violin plots showing median and two quartiles of log10 fold change. Left panels, comparing input counts to n=4 sample counts on d27+d48. Right panels, comparing n=4 counts on d27+d48 to n=4 counts after 21 days of 4 nM vincristine exposure in the presence of stromal support. n=10 sgRNAs for the selected genes and the neutral control LacZ (green); n=2 for essential control genes MYC or PRL9 (red). # indicates genes designated as ‘common essential’ in DepMaP (**Table S2 tab T8**). These are genes for which 90% of cell lines rank the particular gene above a given dependency cutoff. Note: results shown in Fig. 5 and Fig. 6 were obtained by a contemporaneous transduction of all 97 targeted genes but are split over several graphs for display purposes.

**Figure 6.**
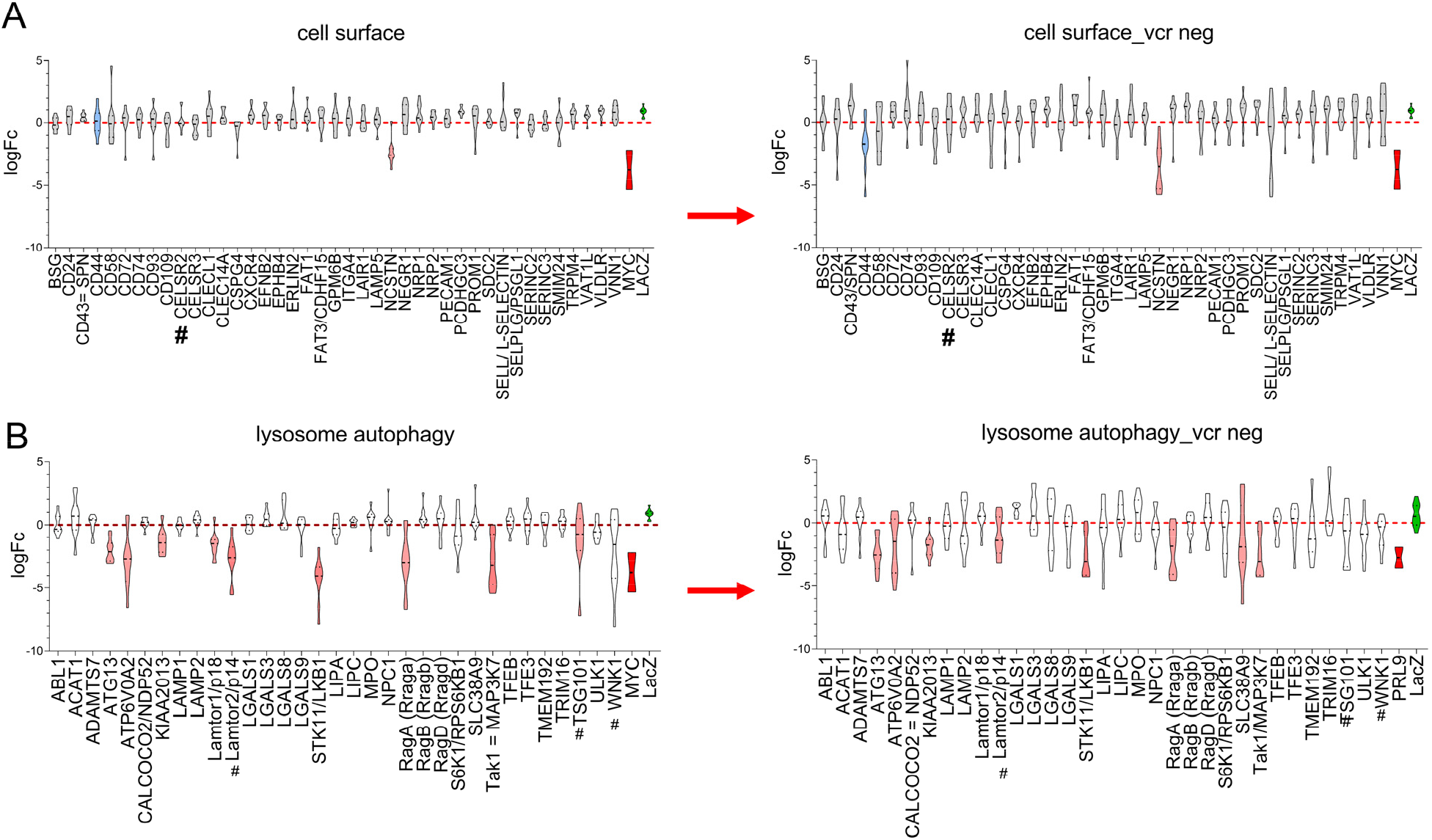
Contribution of selected genes to overall survival and drug-tolerant persister state of KOPN1 BCP-ALL cells. Violin plots showing median and two quartiles of log10 fold change. Left panels, comparing input counts to counts on d27+d48. Right panels, comparing counts on d27+d48 to counts after 21 days of 4 nM vincristine exposure in the presence of stromal support. n=10 sgRNAs for the selected genes and the neutral control LacZ; n=2 for essential control genes MYC or PRL9. **#** indicates genes designated as ‘common essential’ in DepMaP (**Table S2 tab T8**).

As shown in **Figure S6A**, as expected, genes that were included as positive controls needed for survival (boxed red) scored among the most essential in this screen. Drug treatment yielded a further differentiation in the essential functions of some genes. For the positive control genes (**Figure S6B**, boxed red) further loss due to drug treatment was less prominent. Among the target genes, some of the genes flagged as more essential for overall cell survival remained important and vectors expressing their sgRNAs continued to be lost over the period of 21 days of drug treatment.

The Cas9/CRISPR dropout screen identified MYH9 as a gene important for overall survival of these cells (**Figure 5A,** left panel) and for continued survival of DTP cells (**Figure 5A**, right panel). ATP1B3, the beta-3 subunit of a sodium/potassium transporting ATPase which can also transport the chemotherapy drug cisplatin (32), and SLC7A5, which forms a heterodimer with SLC3A2, to transport large neutral amino acids, contributed to overall cell survival (**Figure 5B** left panel). Interestingly, ABCC1 function was dispensable for steady-state growth but became essential for survival of BCP-ALL cells treated with vincristine (**Figure 5B**, compare left and right panels). Of the cell surface proteins assayed here, only NCSTN appeared essential (**Figure 6A**, left), whereas CD44 and CD58 additionally contributed to fitness of DTP cells (**Figure 6A**, right). Finally, many of the genes grouped together in the category lysosomal function and autophagy promoted KOPN1 steady-state fitness and contributed to drug-exposure survival (**Figure 6B**, left). Although Lamtor1 sgRNAs were underrepresented in KOPN1 cells growing under steady-state conditions, there was no further negative selection after treatment with vincristine (**Figure 6B**, right). In contrast, SLC38A9, an mTORC1 activator and arginine sensor (33), was not needed for normal growth but contributed to fitness of DTP BCP-ALL cells (**Figure 6B**).

## DISCUSSION

### Genes with common differential expression in DTP and MRD BCP-ALL cells

In this study we modelled the emergence of DTP cells that survive treatment with the common chemotherapy drug vincristine. A comparison of our data set with other analysis of candidate DTP BCP-ALL cells revealed that some genes were universally increased across the different types of cells, including *in vitro* co-culture, PDX-derived cells and primary patient samples. Notably this included genes that regulate cell motility and adhesion such as EMP1, CXCL16, CYTH4, OSBPL3, GSN and RhoC. Seeing that vincristine is known to increase amoeboid-type cell motility (34), increased expression of such genes could be a chemotherapy-specific feature of the LTP leukemia cells. OSBPL3, which is related to ORP1L, additionally could be involved in lysosomal membrane repair through interactions with VAPA (35) subsequent to vincristine-generated lysosomal membrane damage (29).

Elevated EMP1 is associated with prednisole resistance and worse outcome in BCP-ALL (36, 37). Interestingly, its expression is increased upon adhesion to stromal cells. Moreover it promotes migration and adhesion to stromal cells (36). GSN is an actin-binding, Ca^2+^-regulated protein involved in cell motility and cell shape (38). Increased GSN expression has previously also been linked to vincristine-resistance in a xenograft model of childhood ALL (39). Gelsolin moreover protects cells against apoptosis (40, 41). Although a comparison of GSN RNA expression in 10 matched diagnosis [untreated] and relapsed samples (23) did not show increased expression in samples of relapsed patients, who are presumed to have received prior chemotherapy (**Figure S7A**), our previous multi-omics studies comparing primary B-cell precursor mixed-lineage leukemia (MLL-r) patient cells with normal bone marrow controls measured significant upregulation of both GSN protein and transcript in the MLL-r cells (13). This suggests that increased GSN expression may be an overall hallmark of malignant transformation of normal B-cell precursors. As shown in **Figure S7B**, analysis of gene expression data of pediatric BCP-ALL diagnosis samples supports increased GSN expression in subsets of leukemia including the MLL-r subtype.

There were fewer LTP/MRD genes with common differential expression related to regulation of DNA and RNA. The increased expression of the CDKN1A gene is consistent its regulation by p53 and its well-known activity of inhibiting cell cycle progression at G1. MXD4 is a negative regulator of MYC (42) and its increased expression is consistent with GSEA showing downregulation of MYC pathways (**Table S1 Tab T9**). The DNA methyltransferase PRDM8 is needed for hematopoietic differentiation from embryonic stem cells (43, 44) but no involvement in leukemia has been reported. METTL7A, a member of a family that m6A methylates RNA, was linked to drug resistance (45) and promoting cell survival under glucose deprivation (46) in other cell types.

### Genes needed for overall fitness

We also tested a selected cohort of genes for a possible contribution to the drug-tolerant state when BCP-ALL cells were co-cultured with stromal support. The secreted connective tissue growth factor CTGF/CNN2 is highly overexpressed in BCP-ALL cells (47) and reduction in levels using shRNA was reported to reduce growth and promoted apoptosis (48). However, here it was found not to affect cell survival even under conditions of chemotherapy treatment. The dependence on MYH9 found here was of interest. MYH9 is an ATP-driven motor protein that regulates actomyosin contraction of cells, cell migration and morphology (49). We found that chemotaxis of ALL cells toward supportive stroma and SDF-1α is, in part, regulated by MYH9, as treatment of ALL cells with the specific MYH9 inhibitor blebbistatin significantly decreased their adhesion and migration (50). MYH9 was detected as binding to GSN in a high throughput screen (51). It is needed for survival of HSPC (52), regulates mature B-cell functions (53) and has an important contribution in hematologic malignancies (54–56).

SLC7A5 and SLC3A2 form, among others, a plasma membrane leucine / glutamine antiporter (57). SLC7A5 was shown to regulate survival of T-ALL (58, 59) whereas SLC3A2 is important in B-ALL and AML (60, 61). SLC7A5 and SLC3A2 are also present on lysosomal membranes, promoting leucine uptake into the lysosomal lumen (62). Interestingly, other genes involved in lysosomal function and autophagy such as ATP1B3, LAMTOR2, TSG101 and WNK1 also played an important role in fitness of BCP-ALL cells, similar to their general contributions to cancer fitness (**Table S2 Tab T8**). Remarkably, based on lack of embryonic lethality in mice, an overall loss of function of these genes (RagA, RagB, RagD, TFEB, ATG13, TRIM16, Calcoco2, STK11, S6K1/RPS6KB1, ULK1 ACAT1) is tolerated at an organismal level (Table S21 in (63)) indicating a specific dependence on them of cancer cells. STK11/LKB1 appeared more specifically essential for hematologic malignancies (64) in agreement with our findings here for BCP-ALL survival.

We also tested KIAA2013, a gene of unknown function, based on increased protein levels in a relapse but not diagnosis primary BCP-ALL sample (13) compared to normal CD19+CD10+ pre-B cells. KIAA2013 is a putative single pass type I membrane protein (65) that contains a signal peptide sequence (66). Its ablation was reported to reduce proliferation in among others a CML, an AML and two Multiple Myeloma cell lines (67). Its presence in the lysosomal proteome [table S2 in (66)] indicates that part of its function is linked to this organelle or processes associated with it. KIAA2013 expression correlates with that of the CB2/CNR2 receptor (68) which was identified as a possible treatment target in BCP-ALL (64) and according to Biogrid, it interacts with lysosomal-associated proteins including ATP6V0A2 and cell surface proteins (ATP1B3, NCSTN).

Few of the cell surface protein-encoding genes tested here were needed for survival, indicating that the increased expression noted in many studies correlates with, but is not essential to maintaining leukemia cell homeostasis. CELSR2 appears to be more critically important overall in cancer types than in BCP-ALL co-cultured with stromal cells where its ablation essentially had no effect. Interestingly, in a recent study CELSR2 was shown to contribute to glucocorticoid resistance of BCP-ALL cells (69).

We here found that NCSTN function is needed for BCP-ALL survival. NCSTN is a non-catalytic subunit of the gamma secretase complex that proteolytically cleaves numerous membrane-bound substrates (70) including Notch. Null mutant NSCTN mice die at around E10 (71). Using Vav1-driven knockout of NCSTN in hematopoietic stem cells, Klinakis et al (72) showed that NCSTN loss results in chronic myelomonocytic leukemia in mice. It also contributes to development of mature B-1 B cells (73). Since the critical pathway affected by NCSTN ablation could be Notch signalling, our studies point to NCSTN as an additional gene besides ABCC1 that could be targeted to eliminate DTP BCP-ALL cells treated with standard induction chemotherapy. Interestingly, the importance of NOTCH signalling in BCP-ALL was recently shown experimentally by Fernández-Sevilla et al (74) and Kamga et al (75).

### Genes of importance for DTP development

We identified CD44 and ABCC1 [MRP1] as genes of which loss-of-function did not affect steady-state growth, but did inhibit the ability of the BCP-ALL cells to survive a 21-day vincristine treatment. CD44 binds to hyaluronic acid in the bone marrow extracellular matrix and is an MRD marker in BCP-ALL (76). In AML it increased resistance to a BCL2 inhibitor by promoting SDF1α-CXCR4 interactions (77). In T-ALL, CD44 increased doxorubicin resistance by promoting the increased efflux of this drug through an unknown mechanism. (78). Thus CD44-mediated protection may be mediated though the interactions of the BCP-ALL cells with the microenvironment or functions associated with motility and adhesion of the BCP-ALL cells to stroma.

Drug transporters associated with multi-drug resistance (MDR) in cancer include members of the ABC family of transporters. Specific efflux of vincristine was reported for ABCC1/MRP1, ABCC2/MRP2, ABCC10 and ABCB5/ABCC1/P-gp in different cancer cell types (27). However, in BCP-ALL, overexpression of MDR genes is not considered a major mechanism of drug resistance although higher ABCA2, ABCA3, ABCB1/P-glycoprotein and ABCC1/MRP1 expression was reported (79, 80) with a second publication (81) noting high expression of ABCC2, ABCC3, ABCC4 and ABCC6 with low expression of ABCC1 in relapsed pediatric BCP-ALL patients. Our results show that expression levels do not necessarily equate with functional importance. We did not find evidence that the transcription of the transporters examined here was highly increased in ICN13 BCP-ALL cells treated with chemotherapy [**Table S2 tab T6**]. However, ABCC1/MRP1 was clearly critical for the development of the DTP state of KOPN1 cells treated with vincristine. Although other drug transporters could have compensated for loss of ABCC1 function, this did not appear to happen here within the 21-day treatment period. This could be because other known vincristine transporters [ABCC2, ABCB5, ACBC10-(27)] had very low expression in these cells [**Table S2 tab T6**]. Robey et al (82) indicated that interest in developing inhibitors for MDR genes has decreased. More recently, a new approach reported compounds that could inhibit ABCC1 (83), suggesting it may be possible to inhibit ABCC1 as a strategy to enhance the effects of induction chemotherapy that includes vincristine and eliminate MRD/LTP cells.

### Common features of DTP BCP-ALL

Based on transcriptomics, different mechanisms have been proposed over time to explain tumor or leukemia cell persistence after drug treatment because of microenvironmental protection. Features identified in leukemia cells remaining after chemotherapy include dormancy and a diapause-like state (14, 84, 85). Based on numerous studies, it seems clear that indeed increased quiescence is a common feature of MRD cells in BCP-ALL as well: Turati et al (86), Ebinger et al (14) and our own studies all showed that MRD/DTP cells have reduced activity of MYC and E2F pathways. The reduced MYC pathway activity could be additionally related to the ability of MYC to downregulate lysosomal biogenesis and autophagy (87): our finding that many proteins involved in lysosomal homeostasis and autophagy are needed for general survival as well as that of drug-exposed cells suggests there is an ongoing need to clear such damage.

The increased characteristics of an inflammatory response seen in our study are consistent with the presumed ongoing cellular senescence and cell death in the population as a whole under such circumstances. Interestingly, Turati et al (86), Ebinger et al (14) and our results also noted signatures consistent with increased complement /coagulation and JAK/STAT pathway activation in the MRD BCP-ALL cells. Expression of complement associated genes has been found in carcinomas [reviewed in (88)], and JAK/STAT signalling was recently reported as a hallmark of drug-resistant prostate cancer (89). Thus, increased inflammation is a second common hallmark of MRD/DTP BCP-ALL cells.

Overexpression of genes regulating cell motility and adhesion as well as the loss-of-function effect of MYH9 and CD44 on persistence of drug-treated cells found here are furthermore consistent with a need for the BCP-ALL DTP/MRD cells to interact with the stromal cells and the ECM they produce. These results also are concordant with Ebinger et al (14) who found MRD-like cells to preferentially localize close to the endosteum, with enrichment in genes associated with cell adhesion. These leukemia-niche interactions provide numerous molecular mechanisms for protection (90) and because of this can also be used to target MRD (17).

The landmark paper by Turati et al (86) used primary patient samples and PDX transplant models to shed new light on the nature of the MRD cells that persist after induction chemotherapy of BCP-ALL. These authors showed that the *genetic* heterogeneity present in input cells is maintained within the population of cells that remain viable after chemotherapy in BCP-ALL. Importantly, they showed there is phenotypic heterogeneity present in the original population before treatment, and that chemotherapy forms a bottleneck through which MDR-like cells are produced with less transcriptional heterogeneity. These conclusions were applicable to BCP-ALL cells with different original driver mutations including Ph-like and normal karyotype, whereas our studies used MLL-r BCP-ALL cells. Thus, it is possible that MRD /DTP cells, regardless of genetic lesion, have a common set of phenotypic characteristics that could be targeted specifically at day 30 of induction chemotherapy to definitively eradicate them and prevent future relapses.

## Abbreviations

BCP-ALL: precursor B-cell acute lymphoblastic leukemia
DE: differential expression
DTP: drug-tolerant persister
EMDR: environment-mediated drug resistance
GSEA: Gene Set Enrichment analysis
logFc: log2-fold change
MAGeCK: Model-based Analysis of Genomewide CRISPR/Cas9 Knockout
MFI: mean fluorescent intensity
MRD: minimal residual disease
rpkm: reads per kilobase million

## Acknowledgements

We thank Sachith Gallolu for GSEA analysis. The SC2 Core and Li Fan at Children’s Hospital Los Angeles are acknowledged for RNA-seq. This study was partly supported in 2016/2017 by a New Idea Award from the Leukemia Lymphoma Society and R01 CA090321 and CA172040 to NH.

## Supplemental data

This article contains supplemental data.

## Data availability

RNA-seq data have been deposited in GEO under number GSE176366.

---

**Supplemental Figure 1.**
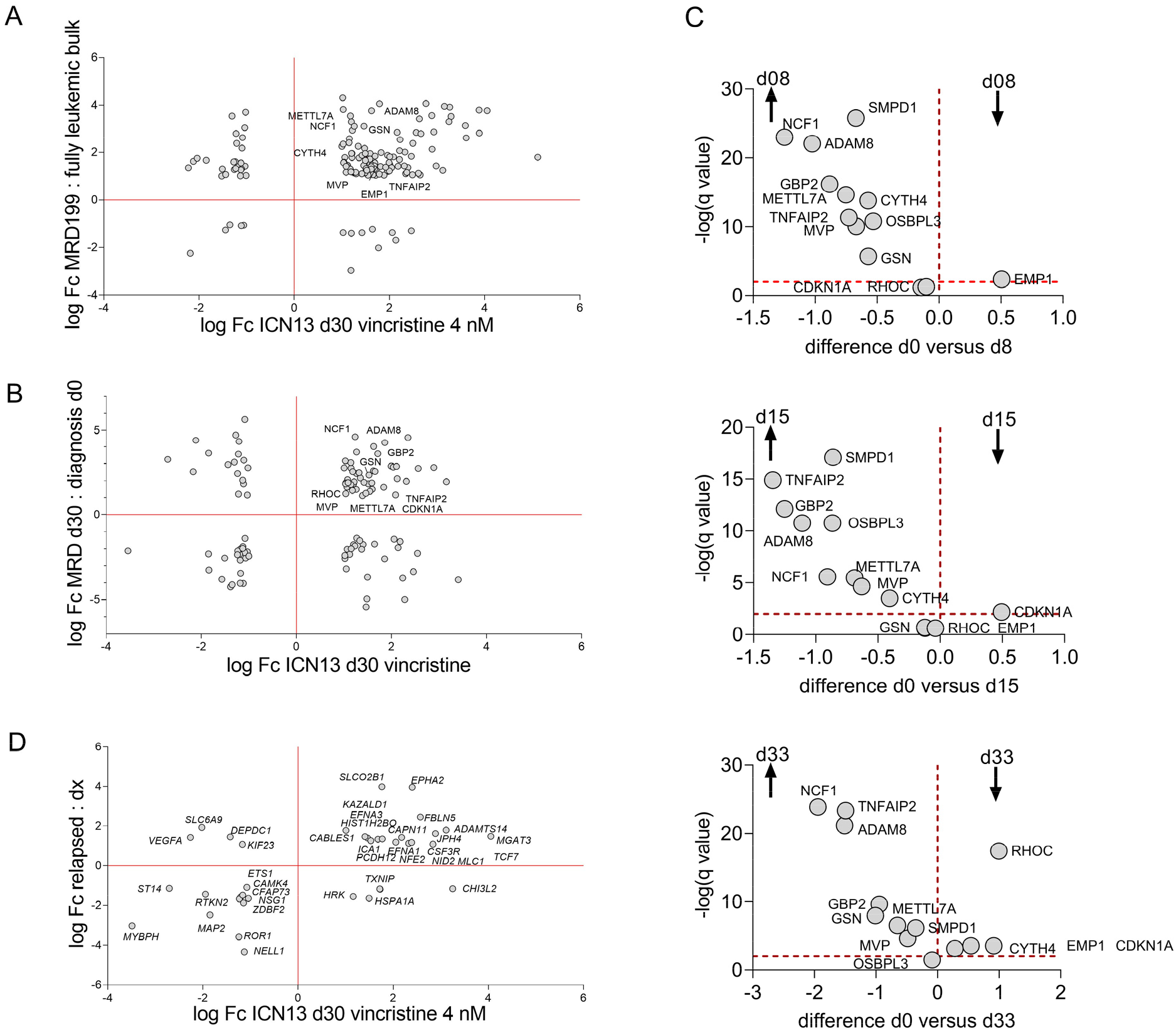
Overlap in significantly differentially expressed genes in ICN13 day 30 vincristine MRD cells. (**A, B**) logFc of DE genes in MRD cells compared to non-chemotherapy treated BCP-ALL cells (GSE83142): (**A**) PDX 199 BCP-ALL transplant model: logFc of DE genes in n=14 purified MRD samples compared to n=8 nonchemotherapy treated BCP-ALL samples. Vincristine/cyclophosphamide, MRD199bulk counts. (**B)** LogFc of DE genes in n=4 diagnosis patient samples and n= 3 samples on day 30 induction chemotherapy. The MDR samples at d30 were purified. (**C**) Volcano plots showing difference in mean gene expression microarray MFI values from GSE67684 (22) in BCP ALL patient samples at day 8 [n= 193, top], d15 [n =49; mid] or d33 [= 59; bottom] of induction chemotherapy compared to values at diagnosis d0 [n = 194]. Dashed line on Y-axis, −log(q value) = 2. Values of single probes per gene are shown. (**D**) LogFc of genes with DE in relapsed (n=10) compared to diagnosis (n=10) samples (23). Dashed line, Y = 2. Not all genes are marked in A, B; see **Table S1 tab T3-T8**.

**Supplemental Figure 2.**
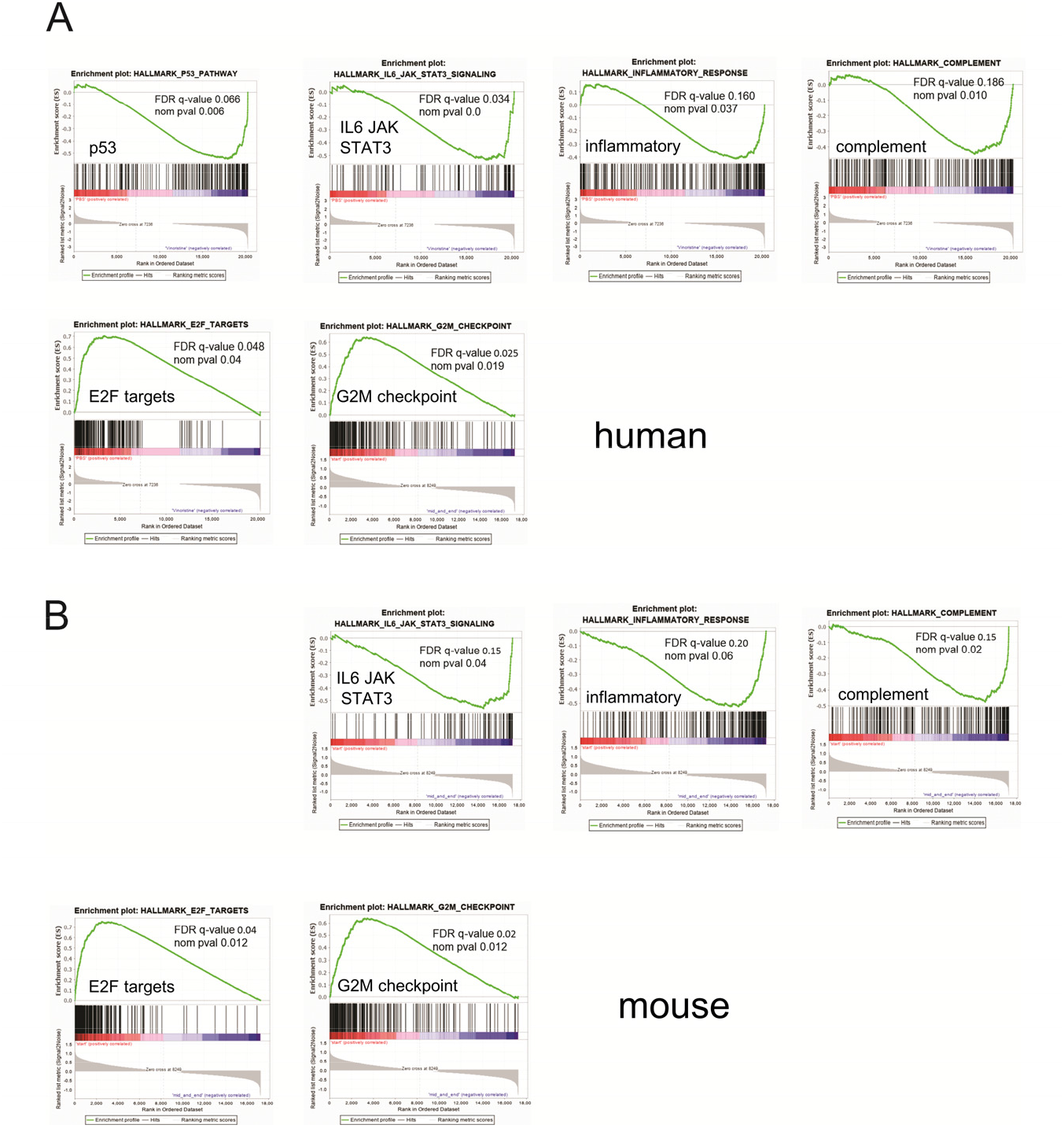
RNA-seq DE results analysed by GSEA (Broad Institute). (**A**) Human ICN13 BCP-ALL cells treated for 30 days with 4 nM vincristine compared to d30 cells treated with PBS. Analysis included 20,287 protein-encoding genes. The amount of gene sets enriched in PBS controls was 19 [of which 2 at FDR<25%; 0 at pval <1%, 2 at pval <5%]; the number of gene sets enriched in vincristine-treated cells was 31 [of which 10 at FDR <25%; 4 at pval <0.1%; 8 at pval <5%]. Also see **Table S1 Tab 5.** (**B**) GSEA of GSE37286 (20) microarray gene expression on murine Bcr/Abl BCP-ALL 8093 or B2 cells treated with 20-50 nM nilotinib or with 1–0.25 mM lonafarnib. Analysis included combined results at t=0 (no drug, n=12 samples) to expression at the mid + endpoint (between d20 and d30) of drug treatment (n =22 samples). The number of gene sets enriched in samples at t=0 compared to mid+end was 20; the number of gene sets enriched in the mid/end phase of drug treatment was 30. Also see **Table S1, Tab 9-10**.

**Supplemental Figure 3.**
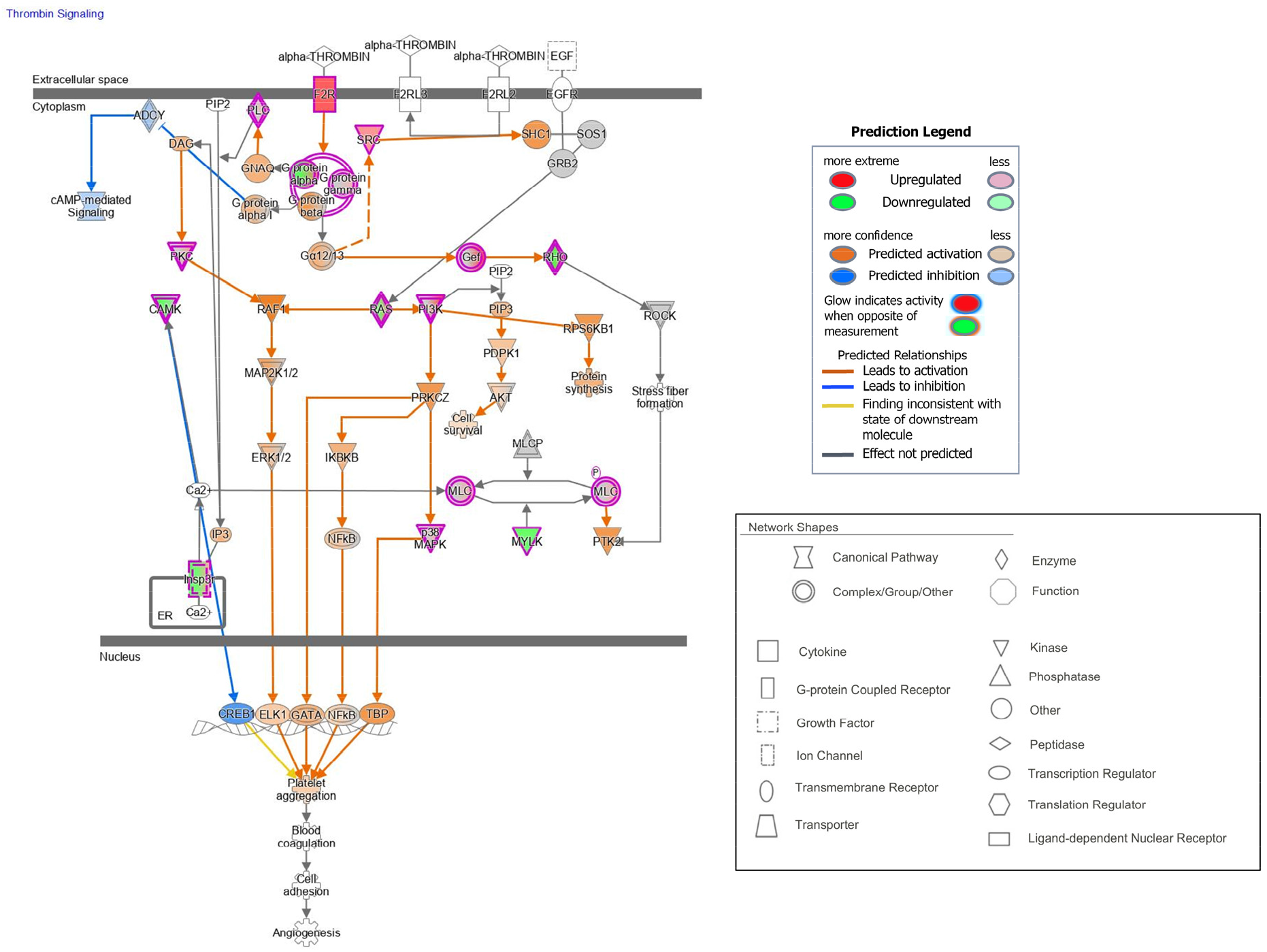
Thrombin signal transduction pathway. IPA analysis indicating involvement of genes significantly regulated in ICN13 BCP-ALL cells treated for 30 days with 4 nM vincristine compared to controls. Keys to colour legend and the network shapes are indicated to the right.

**Supplemental Figure 4.**
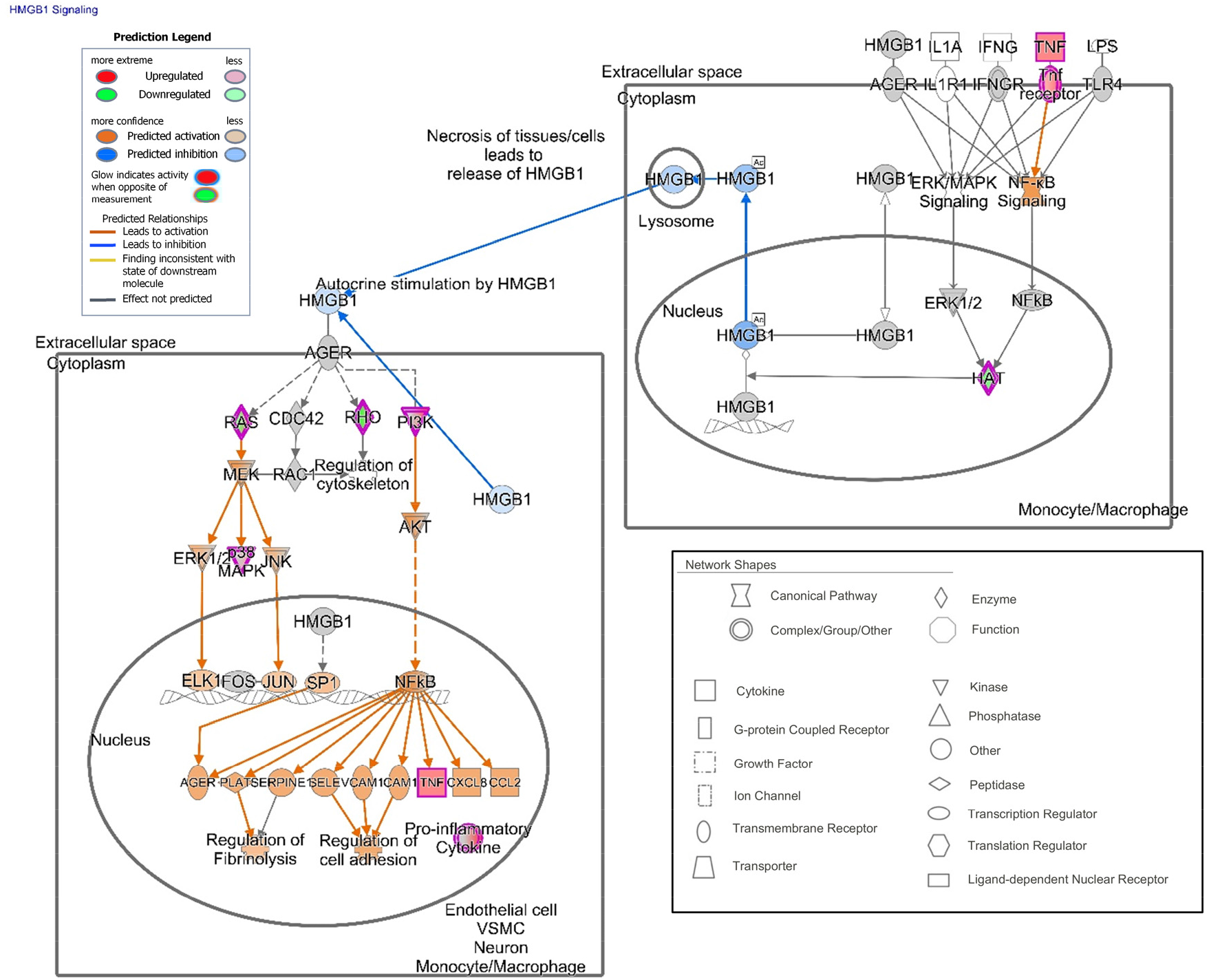
Genes related to the HMBG1 pathway with differential expression in ICN13 at day 30 in 4 nM vincristine-treated BCP-ALL cells. IPA analysis indicating involvement of genes belonging to the HMBG1 pathway significantly regulated in ICN13 BCP-ALL cells treated for 30 days with 4 nM vincristine compared to controls.

**Supplemental Figure 5.**
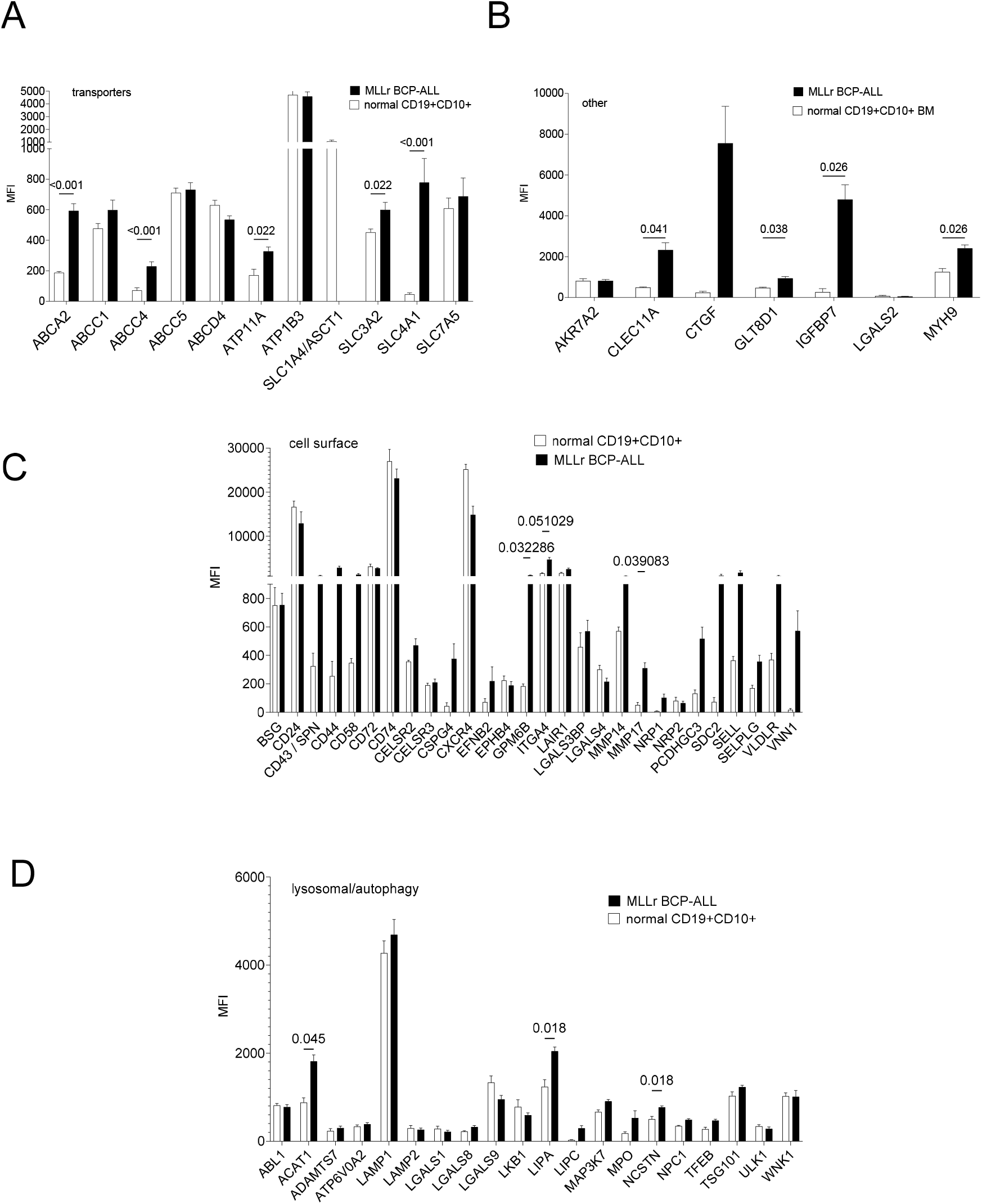
Differential expression of genes selected in Cas9/CRISPR drop-out screen in normal control CD19+CD10+ bone marrow cells compared to primary BCP-ALL samples. Gene expression by microarray reported in GSE28497 (76). MFI, mean fluorescence intensity; n =4 normal pre-B samples; n =17 MLLr-BCP-ALL samples. Unpaired t-tests, exact p-values if <0.05 are shown. Two-stage step-up (Benjamini, Krieger, and Yekutieli). Classes include (**A**) Transporters. ABCC1, ABCC4 and ABCC5 are also known as MRP1, MRP4 and MRP5. (**B**) other genes with DE (reported in (91) table S13 among the top 100 specific for MLL-r subclass of leukemia) and in GSE176364 comparing MLL-r BCP-ALL cells to normal pre-B bone marrow cells (13) (**C)** cell surface proteins (**D**) genes involved in lysosomes and autophagy

**Supplemental Figure 6.**
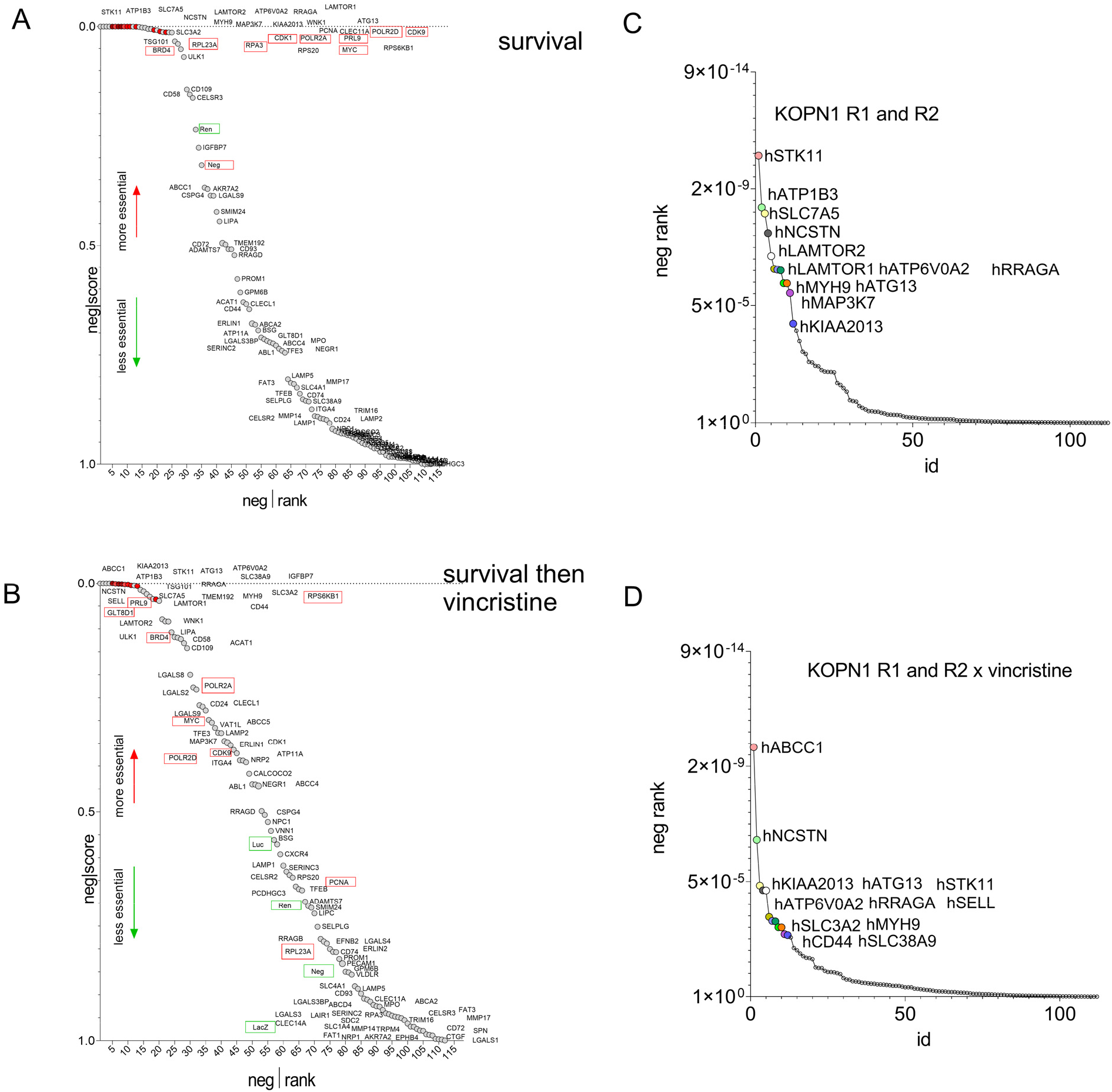
CRISPR-Cas9 dropout screen to identify genes needed for development of drug-permissive survival of BCP-ALL cells treated with vincristine and protected by stromal cells. **(A, B)** Negative-ranked MAGeCK score of selected genes compared to genes with neutral selection activity (green boxes, n=10 sgRNAs/gene) and genes that are essential for cell survival (red boxes, n=2 sgRNAs/gene). **(A**) Comparing input to survival under steady-state conditions. For exact ranking see **Table S3 tab T7**. (B) Comparison of cells containing sgRNAs surviving at d27+d48 with those left after 21-day vincristine treatment. Exact ranking, see **Table S3 tab T8.** (**C, D**). Genes in this screen with the highest negative rank (**Table S3 tab T9**).

**Supplemental Figure 7.**
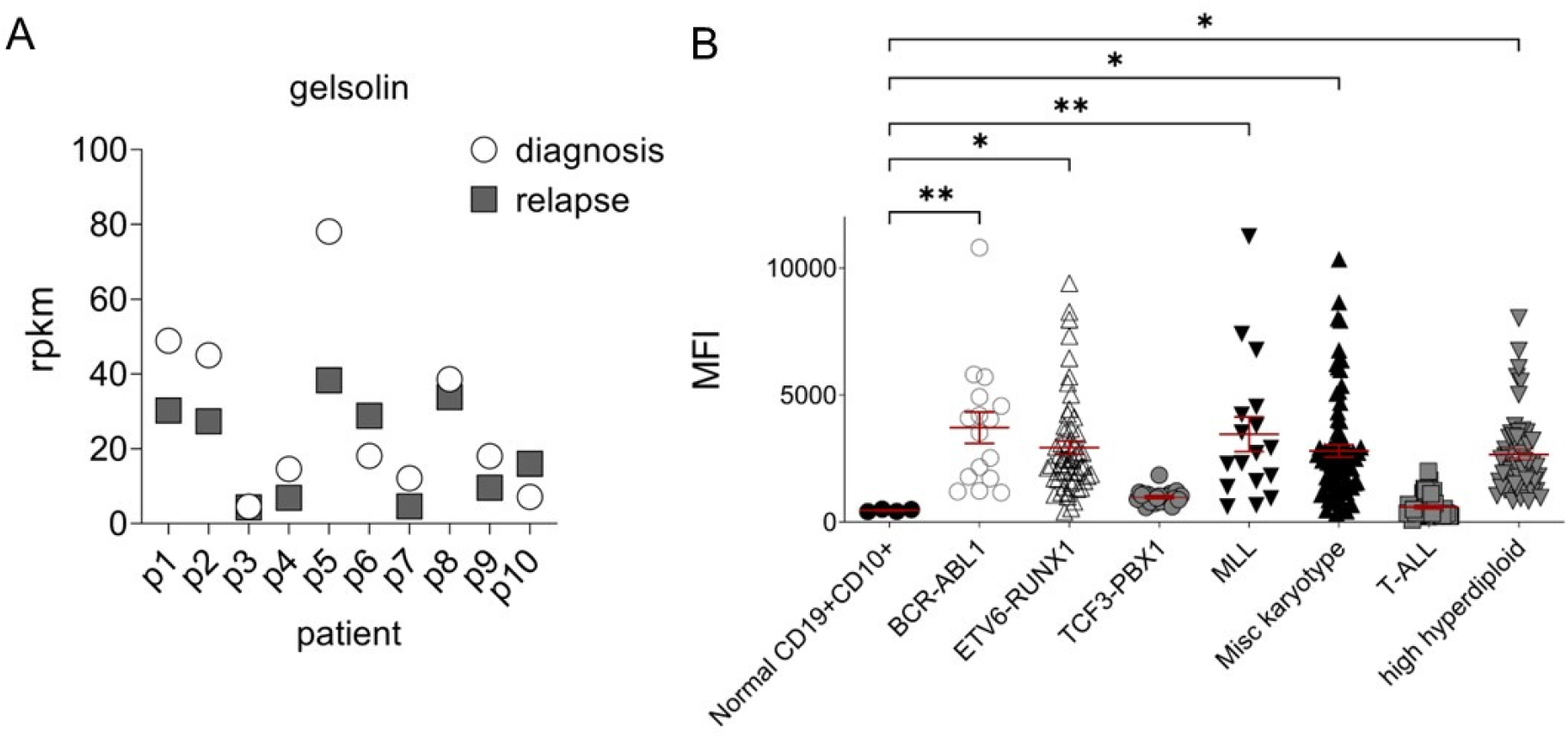
Gelsolin expression is increased in BCP-ALL compared to normal pre-B cells. (**A**) paired diagnosis and relapse BCP-ALL samples (23). (**B**) RNA expression microarray data GSE28497 (76). MFI, mean fluorescence intensity; one way ANOVA, Dunnett’s multiple comparison test, normal pre-B cells to leukemic pre-B cell subtypes. *p<0.05; **p<0.01

## Supplementary Tables

### Table 1 ICN13 MLL-r BCP-ALL cells treated with vincristine

Tab 1 RNAseq results of 18-day treatment 2 nM with vincristine compared to 18-day PBS treated cells.

Tab 2 RNAseq results of 30-day treatment 4 nM with vincristine compared to 30-day PBS treated cells.

Tab 3 Combined RNAseq results of 18-day and 30-day treatment with vincristine compared to PBS treated cells.

Tab 4 Common genes with differential expression in ICN13 on d30 of vcr treatment compared to actively growing d30 PBS treated samples and BCP-ALL MRD cells isolated from PDX mice treated with vincristine plus cyclophosphamide for 14 days compared to non-treated, fully leukemic mice (GSE83142 MRD199bulk).

Tab 5 Overlap of genes with differential expression in ICN13 on d30 of vcr treatment compared to actively growing d30 PBS treated samples and BCP-ALL MRD samples at d30 of induction chemotherapy compared to fully leukemic d0 diagnosis samples (GSE83142 MRDpatients).

Tab 6. Commonly upregulated genes in BCP-ALL cells remaining after long-term chemotherapy treatment.

Tab 7. Expression of commonly upregulated genes (Tab 6) in unfractionated BCP-ALL patient samples at d0, d8, d15 and d33 of induction chemotherapy (GSE67684).

Tab 8 Overlap of genes with differential expression in ICN13 [d30 of vcr treatment compared to actively growing d30 PBS treated samples] and BCP-ALL samples at relapse compared to untreated diagnosis from Meyer et al [PMID 23377183]

Tab 9. GSEA of ICN13 d30 vcr compared to d30 control.

Tab10. GSEA of GSE37286 murine 8093 and B1 Bcr/Abl-expressing pre-B ALL cells treated with nilotinib or lonafarnib

Tab 11. IPA canonical pathways ICN13x vincristine d30

Tab 12. IPA p53 pathways ICN13x vincristine d30

Tab 13 IPA integrin signalling ICN13x vincristine d30

Tab 14 IPA HMBG1 signalling ICN13x vincristine d30

Tab 15. IPA thrombin signalling ICN13x vincristine d30

Tab 16 IPA kinetochore metaphase ICN13x vincristine d30

Tab 17 IPA G2M checkpoint ICN13x vincristine d30

Tab 18 IPA inflammatory response ICN13x vincristine d30

Tab 19 IPA CSF2 pathway ICN13x vincristine d30

Tab 20 IPA SATB1 pathway ICN13x vincristine d30

Tab 21. Genes with DE that belong to autophagy cohort listed in PMID 28552616 (66)

Tab 22. Genes with DE that belong to stressome described in PMID 32105731 (92)

Tab 23. Genes with DE involved in autophagy described in PMID 29700228 (93)

### Table 2. Characteristics of 97 genes tested in Cas9/CRISPR drop-out assay for effect on KOPN1 MLL-r BCP-ALL cell survival and drug resistance against vincristine

Tab 1. Genes differentially expressed in ICN13 BCP-ALL cells treated with 4 nM vincristine for 30 days [Table S1].

Tab 2. Genes with differential expression in MRD samples from patients compared to diagnosis (GSE83142 MRDpatients).

Tab 3. Genes with differential expression in PDX-isolated MRD199bulk cells (GSE83142 MRD199bulk)

Tab 4. Genes showing differential expression in primary samples of pediatric BCP-ALL treated with induction chemotherapy d0 compared to day 15 and day 33 (GSE67684).

Tab 5. Tested genes with different expressed in primary MLL-r BCP-ALL patient samples (n=3) compared to normal bone marrow CD19+CD10+ controls (n=4) (GSE176364).

Tab 6. ABC transporter expression in BCP-ALL; left GSE176364; right, Table S1

Tab 7. List of genes and Uniprot links

Tab 8. Essentiality in SANGER Cancer Dependency Map https://depmap.sanger.ac.uk/ and https://depmap.org/portal/

### Table 3. DNA counts and MAGeCK analysis of 97-selected gene screen

Tab 1. Summary counts

Tab 2. Counts

Tab 3. Normalized counts

Tab 4. Normalized counts sorted by alphabetical gene names

Tab 5. Figure 6 and 7 left panels. Analysis (3)sgRNA comparing 0615_plasmid_S33 with [02_S103 + 06_S107 + 04_S105 + 08_S109]>

Tab 6. Figure 6 and 7 right panels. Analysis (6)sgRNA comparing [02_S103 + 06_S107 + 04_S105 + 08_S109] with [14_S115 + 18_S119 + 16_S117 + 20_S121]

Tab 7. Supplemental Figure 6A, C. MAGeCK results genes from comparison in Tab 5, input versus survival on d27+d48.

Tab 8. Supplemental Figure 6B, D. MEGeCK results genes from comparison in Tab 6, survival d27+d48 versus survival after 1 nM vincristine for 21 days.

Tab 9. Summary

## References

1. Terwilliger T, Abdul-Hay M. Acute lymphoblastic leukemia: a comprehensive review and 2017 update. Blood Cancer J. 2017;7(6):e577.

2. Bhm JW, Sia KCS, Jones C, Evans K, Mariana A, Pang I, et al. Combination efficacy of ruxolitinib with standard-of-care drugs in CRLF2-rearranged Ph-like acute lymphoblastic leukemia. Leukemia. 2021.

3. Risinger AL, Giles FJ, Mooberry SL. Microtubule dynamics as a target in oncology. Cancer Treat Rev. 2009;35(3):255–61.

4. Contreras Yametti GP, Ostrow TH, Jasinski S, Raetz EA, Carroll WL, Evensen NA. Minimal Residual Disease in Acute Lymphoblastic Leukemia: Current Practice and Future Directions. Cancers (Basel). 2021;13(8).

5. Nguyen K, Devidas M, Cheng SC, La M, Raetz EA, Carroll WL, et al. Factors influencing survival after relapse from acute lymphoblastic leukemia: a Children’s Oncology Group study. Leukemia. 2008;22(12):2142–50.

6. Meads MB, Gatenby RA, Dalton WS. Environment-mediated drug resistance: a major contributor to minimal residual disease. Nat Rev Cancer. 2009;9(9):665–74.

7. Bakker E, Qattan M, Mutti L, Demonacos C, Krstic-Demonacos M. The role of microenvironment and immunity in drug response in leukemia. Biochim Biophys Acta. 2016;1863(3):414–26.

8. Fei F, Abdel-Azim H, Lim M, Arutyunyan A, von Itzstein M, Groffen J, et al. Galectin-3 in pre-B acute lymphoblastic leukemia. Leukemia. 2013;27(12):2385–8.

9. Fei F, Joo EJ, Tarighat SS, Schiffer I, Paz H, Fabbri M, et al. B-cell precursor acute lymphoblastic leukemia and stromal cells communicate through Galectin-3. Oncotarget. 2015;6(13):11378–94.

10. Paz H, Joo EJ, Chou CH, Fei F, Mayo KH, Abdel-Azim H, et al. Treatment of B-cell precursor acute lymphoblastic leukemia with the Galectin-1 inhibitor PTX008. J Exp Clin Cancer Res. 2018;37(1):67.

11. Geng H, Hurtz C, Lenz KB, Chen Z, Baumjohann D, Thompson S, et al. Self-enforcing feedback activation between BCL6 and pre-B cell receptor signaling defines a distinct subtype of acute lymphoblastic leukemia. Cancer Cell. 2015;27(3):409–25.

12. Chen Z, Shojaee S, Buchner M, Geng H, Lee JW, Klemm L, et al. Signalling thresholds and negative B-cell selection in acute lymphoblastic leukaemia. Nature. 2015;521(7552):357–61.

13. Oliveira T, Zhang M, Joo EJ, Abdel-Azim H, Chen CW, Yang L, et al. Glycoproteome remodeling in MLL-rearranged B-cell precursor acute lymphoblastic leukemia. Theranostics. 2021;11(19):9519–37.

14. Ebinger S, Ozdemir EZ, Ziegenhain C, Tiedt S, Castro Alves C, Grunert M, et al. Characterization of Rare, Dormant, and Therapy-Resistant Cells in Acute Lymphoblastic Leukemia. Cancer Cell. 2016;30(6):849–62.

15. Li W, Koster J, Xu H, Chen CH, Xiao T, Liu JS, et al. Quality control, modeling, and visualization of CRISPR screens with MAGeCK-VISPR. Genome Biol. 2015;16:281.

16. Li W, Xu H, Xiao T, Cong L, Love MI, Zhang F, et al. MAGeCK enables robust identification of essential genes from genome-scale CRISPR/Cas9 knockout screens. Genome Biol. 2014;15(12):554.

17. Hsieh YT, Gang EJ, Geng H, Park E, Huantes S, Chudziak D, et al. Integrin alpha4 blockade sensitizes drug resistant pre-B acute lymphoblastic leukemia to chemotherapy. Blood. 2013;121(10):1814–8.

18. Quagliano A, Gopalakrishnapillai A, Kolb EA, Barwe SP. CD81 knockout promotes chemosensitivity and disrupts in vivo homing and engraftment in acute lymphoblastic leukemia. Blood Adv. 2020;4(18):4393–405.

19. Usmani S, Sivagnanalingam U, Tkachenko O, Nunez L, Shand JC, Mullen CA. Support of acute lymphoblastic leukemia cells by nonmalignant bone marrow stromal cells. Oncol Lett. 2019;17(6):5039–49.

20. Feldhahn N, Arutyunyan A, Stoddart S, Zhang B, Schmidhuber S, Yi SJ, et al. Environment-mediated drug resistance in Bcr/Abl-positive acute lymphoblastic leukemia. Oncoimmunology. 2012;1(5):618–29.

21. Parameswaran R, Lim M, Arutyunyan A, Abdel-Azim H, Hurtz C, Lau K, et al. O-acetylated N-acetylneuraminic acid as a novel target for therapy in human pre-B acute lymphoblastic leukemia. J Exp Med. 2013;210(4):805–19.

22. Yeoh AE, Li Z, Dong D, Lu Y, Jiang N, Trka J, et al. Effective Response Metric: a novel tool to predict relapse in childhood acute lymphoblastic leukaemia using time-series gene expression profiling. Br J Haematol. 2018;181(5):653–63.

23. Meyer JA, Wang J, Hogan LE, Yang JJ, Dandekar S, Patel JP, et al. Relapse-specific mutations in NT5C2 in childhood acute lymphoblastic leukemia. Nat Genet. 2013;45(3):290–4.

24. Subramanian A, Tamayo P, Mootha VK, Mukherjee S, Ebert BL, Gillette MA, et al. Gene set enrichment analysis: a knowledge-based approach for interpreting genome-wide expression profiles. Proc Natl Acad Sci U S A. 2005;102(43):15545–50.

25. Bracken AP, Ciro M, Cocito A, Helin K. E2F target genes: unraveling the biology. Trends Biochem Sci. 2004;29(8):409–17.

26. Fischer M. Mice Are Not Humans: The Case of p53. Trends Cancer. 2021;7(1):12–4.

27. Wu CP, Hsieh CH, Wu YS. The emergence of drug transporter-mediated multidrug resistance to cancer chemotherapy. Mol Pharm. 2011;8(6):1996–2011.

28. de Klerk DJ, Honeywell RJ, Jansen G, Peters GJ. Transporter and Lysosomal Mediated (Multi)drug Resistance to Tyrosine Kinase Inhibitors and Potential Strategies to Overcome Resistance. Cancers (Basel). 2018;10(12).

29. Groth-Pedersen L, Ostenfeld MS, Hoyer-Hansen M, Nylandsted J, Jaattela M. Vincristine induces dramatic lysosomal changes and sensitizes cancer cells to lysosome-destabilizing siramesine. Cancer Res. 2007;67(5):2217–25.

30. Sehgal AR, Konig H, Johnson DE, Tang D, Amaravadi RK, Boyiadzis M, et al. You eat what you are: autophagy inhibition as a therapeutic strategy in leukemia. Leukemia. 2015;29(3):517–25.

31. Smith AG, Macleod KF. Autophagy, cancer stem cells and drug resistance. J Pathol. 2019;247(5):708–18.

32. Khoo XH, Paterson IC, Goh BH, Lee WL. Cisplatin-Resistance in Oral Squamous Cell Carcinoma: Regulation by Tumor Cell-Derived Extracellular Vesicles. Cancers (Basel). 2019;11(8).

33. Wang S, Tsun ZY, Wolfson RL, Shen K, Wyant GA, Plovanich ME, et al. Metabolism. Lysosomal amino acid transporter SLC38A9 signals arginine sufficiency to mTORC1. Science. 2015;347(6218):188–94.

34. Eitaki M, Yamamori T, Meike S, Yasui H, Inanami O. Vincristine enhances amoeboid-like motility via GEF-H1/RhoA/ROCK/Myosin light chain signaling in MKN45 cells. BMC Cancer. 2012;12:469.

35. Radulovic M, Wenzel EM, Gilani S, Holland LK, Lystad AH, Phuyal S, et al. Cholesterol transfer via endoplasmic reticulum contacts mediates lysosome damage repair. EMBO J. 2022;41(24):e112677.

36. Aries IM, Jerchel IS, van den Dungen RE, van den Berk LC, Boer JM, Horstmann MA, et al. EMP1, a novel poor prognostic factor in pediatric leukemia regulates prednisolone resistance, cell proliferation, migration and adhesion. Leukemia. 2014;28(9):1828–37.

37. Singh J, Kumari S, Arora M, Verma D, Palanichamy JK, Kumar R, et al. Prognostic Relevance of Expression of EMP1, CASP1, and NLRP3 Genes in Pediatric B-Lineage Acute Lymphoblastic Leukemia. Front Oncol. 2021;11:606370.

38. Feldt J, Schicht M, Garreis F, Welss J, Schneider UW, Paulsen F. Structure, regulation and related diseases of the actin-binding protein gelsolin. Expert Rev Mol Med. 2019;20:e7.

39. Verrills NM, Liem NL, Liaw TY, Hood BD, Lock RB, Kavallaris M. Proteomic analysis reveals a novel role for the actin cytoskeleton in vincristine resistant childhood leukemia--an in vivo study. Proteomics. 2006;6(5):1681–94.

40. Illescas M, Penas A, Arenas J, Martin MA, Ugalde C. Regulation of Mitochondrial Function by the Actin Cytoskeleton. Front Cell Dev Biol. 2021;9:795838.

41. Kusano H, Shimizu S, Koya RC, Fujita H, Kamada S, Kuzumaki N, et al. Human gelsolin prevents apoptosis by inhibiting apoptotic mitochondrial changes via closing VDAC. Oncogene. 2000;19(42):4807–14.

42. Heckler M, Ali LR, Clancy-Thompson E, Qiang L, Ventre KS, Lenehan P, et al. Inhibition of CDK4/6 Promotes CD8 T-cell Memory Formation. Cancer Discov. 2021;11(10):2564–81.

43. Cypris O, Eipel M, Franzen J, Rosseler C, Tharmapalan V, Kuo CC, et al. PRDM8 reveals aberrant DNA methylation in aging syndromes and is relevant for hematopoietic and neuronal differentiation. Clin Epigenetics. 2020;12(1):125.

44. Di Tullio F, Schwarz M, Zorgati H, Mzoughi S, Guccione E. The duality of PRDM proteins: epigenetic and structural perspectives. FEBS J. 2022;289(5):1256–75.

45. Jun F, Peng Z, Zhang Y, Shi D. Quantitative proteomic analysis identifies novel regulators of methotrexate resistance in choriocarcinoma. Gynecol Oncol. 2020;157(1):268–79.

46. Lee E, Kim JY, Kim TK, Park SY, Im GI. Methyltransferase-like protein 7A (METTL7A) promotes cell survival and osteogenic differentiation under metabolic stress. Cell Death Discov. 2021;7(1):154.

47. Boag JM, Beesley AH, Firth MJ, Freitas JR, Ford J, Brigstock DR, et al. High expression of connective tissue growth factor in pre-B acute lymphoblastic leukaemia. Br J Haematol. 2007;138(6):740–8.

48. Wells JE, Howlett M, Halse HM, Heng J, Ford J, Cheung LC, et al. High expression of connective tissue growth factor accelerates dissemination of leukaemia. Oncogene. 2016;35(35):4591–600.

49. Conti MA, Adelstein RS. Nonmuscle myosin II moves in new directions. J Cell Sci. 2008;121(Pt 1):11–8.

50. Fei F, Zhang M, Tarighat SS, Joo EJ, Yang L, Heisterkamp N. Galectin-1 and Galectin-3 in B-Cell Precursor Acute Lymphoblastic Leukemia. Int J Mol Sci. 2022;23(22).

51. Hein MY, Hubner NC, Poser I, Cox J, Nagaraj N, Toyoda Y, et al. A human interactome in three quantitative dimensions organized by stoichiometries and abundances. Cell. 2015;163(3):712–23.

52. An Q, Dong Y, Cao Y, Pan X, Xue Y, Zhou Y, et al. Myh9 Plays an Essential Role in the Survival and Maintenance of Hematopoietic Stem/Progenitor Cells. Cells. 2022;11(12).

53. Hoogeboom R, Natkanski EM, Nowosad CR, Malinova D, Menon RP, Casal A, et al. Myosin IIa Promotes Antibody Responses by Regulating B Cell Activation, Acquisition of Antigen, and Proliferation. Cell Rep. 2018;23(8):2342–53.

54. Chang F, Kong SJ, Wang L, Choi BK, Lee H, Kim C, et al. Targeting Actomyosin Contractility Suppresses Malignant Phenotypes of Acute Myeloid Leukemia Cells. Int J Mol Sci. 2020;21(10).

55. Seri M, Cusano R, Gangarossa S, Caridi G, Bordo D, Lo Nigro C, et al. Mutations in MYH9 result in the May-Hegglin anomaly, and Fechtner and Sebastian syndromes. The May-Heggllin/Fechtner Syndrome Consortium. Nat Genet. 2000;26(1):103–5.

56. Wigton EJ, Thompson SB, Long RA, Jacobelli J. Myosin-IIA regulates leukemia engraftment and brain infiltration in a mouse model of acute lymphoblastic leukemia. J Leukoc Biol. 2016;100(1):143–53.

57. Bolomsky A, Young RM. Pathogenic signaling in multiple myeloma. Semin Oncol. 2022;49(1):27–40.

58. Rosilio C, Nebout M, Imbert V, Griessinger E, Neffati Z, Benadiba J, et al. L-type amino-acid transporter 1 (LAT1): a therapeutic target supporting growth and survival of T-cell lymphoblastic lymphoma/T-cell acute lymphoblastic leukemia. Leukemia. 2015;29(6):1253–66.

59. Grzes KM, Swamy M, Hukelmann JL, Emslie E, Sinclair LV, Cantrell DA. Control of amino acid transport coordinates metabolic reprogramming in T-cell malignancy. Leukemia. 2017;31(12):2771–9.

60. Bajaj J, Konuma T, Lytle NK, Kwon HY, Ablack JN, Cantor JM, et al. CD98-Mediated Adhesive Signaling Enables the Establishment and Propagation of Acute Myelogenous Leukemia. Cancer Cell. 2016;30(5):792–805.

61. Boutter J, Huang Y, Marovca B, Vonderheit A, Grotzer MA, Eckert C, et al. Imagebased RNA interference screening reveals an individual dependence of acute lymphoblastic leukemia on stromal cysteine support. Oncotarget. 2014;5(22):11501–12.

62. Yang Y, Bolomsky A, Oellerich T, Chen P, Ceribelli M, Haupl B, et al. Oncogenic RAS commandeers amino acid sensing machinery to aberrantly activate mTORC1 in multiple myeloma. Nat Commun. 2022;13(1):5469.

63. Luck K, Kim DK, Lambourne L, Spirohn K, Begg BE, Bian W, et al. A reference map of the human binary protein interactome. Nature. 2020;580(7803):402–8.

64. Chan LN, Chen Z, Braas D, Lee JW, Xiao G, Geng H, et al. Metabolic gatekeeper function of B-lymphoid transcription factors. Nature. 2017;542(7642):479–83.

65. Ghosh D, Lippert D, Krokhin O, Cortens JP, Wilkins JA. Defining the membrane proteome of NK cells. J Mass Spectrom. 2010;45(1):1–25.

66. Wyant GA, Abu-Remaileh M, Frenkel EM, Laqtom NN, Dharamdasani V, Lewis CA, et al. NUFIP1 is a ribosome receptor for starvation-induced ribophagy. Science. 2018;360(6390):751–8.

67. Summary KDG. [Available from: KIAA2013 DepMap Gene Summary

68. Sharaf A, Mensching L, Keller C, Rading S, Scheffold M, Palkowitsch L, et al. Systematic Affinity Purification Coupled to Mass Spectrometry Identified p62 as Part of the Cannabinoid Receptor CB2 Interactome. Front Mol Neurosci. 2019;12:224.

69. Autry RJ, Paugh SW, Carter R, Shi L, Liu J, Ferguson DC, et al. Integrative genomic analyses reveal mechanisms of glucocorticoid resistance in acute lymphoblastic leukemia. Nat Cancer. 2020;1(3):329–44.

70. Moore G, Annett S, McClements L, Robson T. Top Notch Targeting Strategies in Cancer: A Detailed Overview of Recent Insights and Current Perspectives. Cells. 2020;9(6).

71. Li T, Ma G, Cai H, Price DL, Wong PC. Nicastrin is required for assembly of presenilin/gamma-secretase complexes to mediate Notch signaling and for processing and trafficking of beta-amyloid precursor protein in mammals. J Neurosci. 2003;23(8):3272–7.

72. Klinakis A, Lobry C, Abdel-Wahab O, Oh P, Haeno H, Buonamici S, et al. A novel tumour-suppressor function for the Notch pathway in myeloid leukaemia. Nature. 2011;473(7346):230–3.

73. Choi JH, Han J, Theodoropoulos PC, Zhong X, Wang J, Medler D, et al. Essential requirement for nicastrin in marginal zone and B-1 B cell development. Proc Natl Acad Sci U S A. 2020;117(9):4894–901.

74. Fernandez-Sevilla LM, Valencia J, Flores-Villalobos MA, Gonzalez-Murillo A, Sacedon R, Jimenez E, et al. The choroid plexus stroma constitutes a sanctuary for paediatric B-cell precursor acute lymphoblastic leukaemia in the central nervous system. J Pathol. 2020;252(2):189–200.

75. Takam Kamga P, Dal Collo G, Midolo M, Adamo A, Delfino P, Mercuri A, et al. Inhibition of Notch Signaling Enhances Chemosensitivity in B-cell Precursor Acute Lymphoblastic Leukemia. Cancer Res. 2019;79(3):639–49.

76. Coustan-Smith E, Song G, Clark C, Key L, Liu P, Mehrpooya M, et al. New markers for minimal residual disease detection in acute lymphoblastic leukemia. Blood. 2011;117(23):6267–76.

77. Yu X, Munoz-Sagredo L, Streule K, Muschong P, Bayer E, Walter RJ, et al. CD44 loss of function sensitizes AML cells to the BCL-2 inhibitor venetoclax by decreasing CXCL12-driven survival cues. Blood. 2021;138(12):1067–80.

78. Hoofd C, Wang X, Lam S, Jenkins C, Wood B, Giambra V, et al. CD44 promotes chemoresistance in T-ALL by increased drug efflux. Exp Hematol. 2016;44(3):166–71 e17.

79. Aberuyi N, Rahgozar S, Khosravi Dehaghi Z, Moafi A, Masotti A, Paolini A. The translational expression of ABCA2 and ABCA3 is a strong prognostic biomarker for multidrug resistance in pediatric acute lymphoblastic leukemia. Onco Targets Ther. 2017;10:3373–80.

80. Rahgozar S, Moafi A, Abedi M, Entezar EGM, Moshtaghian J, Ghaedi K, et al. mRNA expression profile of multidrug-resistant genes in acute lymphoblastic leukemia of children, a prognostic value for ABCA3 and ABCA2. Cancer Biol Ther. 2014;15(1):35–41.

81. Mehrvar N, Abolghasemi H, Rezvany MR, Akbari ME, Saberynejad J, Mehrvar A, et al. Characterizing Iranian Pediatric Patients With Relapsed Acute Lymphoblastic Leukemia Through Gene Expression Profiling of Common ATP Binding Cassette Transporters Subfamily C. J Pediatr Hematol Oncol. 2020;42(1):41–5.

82. Robey RW, Pluchino KM, Hall MD, Fojo AT, Bates SE, Gottesman MM. Revisiting the role of ABC transporters in multidrug-resistant cancer. Nat Rev Cancer. 2018;18(7):452–64.

83. Silbermann K, Stefan SM, Elshawadfy R, Namasivayam V, Wiese M. Identification of Thienopyrimidine Scaffold as an Inhibitor of the ABC Transport Protein ABCC1 (MRP1) and Related Transporters Using a Combined Virtual Screening Approach. J Med Chem. 2019;62(9):4383–400.

84. Senft D, Jeremias I. A rare subgroup of leukemia stem cells harbors relapse-inducing potential in acute lymphoblastic leukemia. Exp Hematol. 2019;69:1–10.

85. Dhimolea E, de Matos Simoes R, Kansara D, Al’Khafaji A, Bouyssou J, Weng X, et al. An Embryonic Diapause-like Adaptation with Suppressed Myc Activity Enables Tumor Treatment Persistence. Cancer Cell. 2021;39(2):240–56 e11.

86. Turati VA, Guerra-Assuncao JA, Potter NE, Gupta R, Ecker S, Daneviciute A, et al. Chemotherapy induces canalization of cell state in childhood B-cell precursor acute lymphoblastic leukemia. Nat Cancer. 2021;2(8):835–52.

87. Annunziata I, van de Vlekkert D, Wolf E, Finkelstein D, Neale G, Machado E, et al. MYC competes with MiT/TFE in regulating lysosomal biogenesis and autophagy through an epigenetic rheostat. Nat Commun. 2019;10(1):3623.

88. Roumenina LT, Daugan MV, Petitprez F, Sautes-Fridman C, Fridman WH. Context-dependent roles of complement in cancer. Nat Rev Cancer. 2019;19(12):698–715.

89. Chan JM, Zaidi S, Love JR, Zhao JL, Setty M, Wadosky KM, et al. Lineage plasticity in prostate cancer depends on JAK/STAT inflammatory signaling. Science. 2022;377(6611):1180–91.

90. Meyer LK, Hermiston ML. The bone marrow microenvironment as a mediator of chemoresistance in acute lymphoblastic leukemia. Cancer Drug Resist. 2019;2(4):1164–77.

91. Ross ME, Mahfouz R, Onciu M, Liu HC, Zhou X, Song G, et al. Gene expression profiling of pediatric acute myelogenous leukemia. Blood. 2004;104(12):3679–87.

92. Kuechler ER, Budzynska PM, Bernardini JP, Gsponer J, Mayor T. Distinct Features of Stress Granule Proteins Predict Localization in Membraneless Organelles. J Mol Biol. 2020;432(7):2349–68.

93. Li X, Yu W, Qian X, Xia Y, Zheng Y, Lee JH, et al. Nucleus-Translocated ACSS2 Promotes Gene Transcription for Lysosomal Biogenesis and Autophagy. Mol Cell. 2017;66(5):684–97 e9.

